# TFE3 fusions direct an oncogenic transcriptional program that drives OXPHOS and unveils vulnerabilities in translocation renal cell carcinoma

**DOI:** 10.1101/2024.08.09.607311

**Authors:** Jiao Li, Kaimeng Huang, Fiona McBride, Ananthan Sadagopan, Daniel S Gallant, Meha Thakur, Prateek Khanna, Bingchen Li, Maolin Ge, Cary N. Weiss, Mingkee Achom, Qingru Xu, Kun Huang, Birgitta A. Ryback, Miao Gui, Liron Bar-Peled, Srinivas R. Viswanathan

## Abstract

Translocation renal cell carcinoma (tRCC) is an aggressive subtype of kidney cancer driven by *TFE3* gene fusions, which act via poorly characterized downstream mechanisms. Here we report that TFE3 fusions transcriptionally rewire tRCCs toward oxidative phosphorylation (OXPHOS), contrasting with the highly glycolytic metabolism of most other renal cancers. This TFE3 fusion-driven OXPHOS program, together with heightened glutathione levels found in renal cancers, renders tRCCs sensitive to reductive stress – a metabolic stress state induced by an imbalance of reducing equivalents. Genome-scale CRISPR screening identifies tRCC-selective vulnerabilities linked to this metabolic state, including *EGLN1*, which hydroxylates HIF-1α and targets it for proteolysis. Inhibition of EGLN1 compromises tRCC cell growth by stabilizing HIF-1a and promoting metabolic reprogramming away from OXPHOS, thus representing a vulnerability to OXPHOS-dependent tRCC cells. Our study defines a distinctive tRCC-essential metabolic program driven by TFE3 fusions and nominates EGLN1 inhibition as a therapeutic strategy to counteract fusion-induced metabolic rewiring.

## Main

Translocation renal cell carcinoma (tRCC) comprises 1-5% of renal cell carcinomas (RCCs) in adults and a majority of RCCs in children^1–3^. tRCC is clinically aggressive and lacks effective therapies, therefore representing a major unmet need amongst kidney cancers^4,5^. Genetically, tRCC is driven by an activating fusion involving a transcription factor in the MiT/TFE gene family, most commonly *TFE3*^1,6^. These translocations, which occur between *TFE3* and any one of several different partner genes, result in the expression of a transcription factor fusion protein that is constitutively nuclear^3^.

Therapies that are effective for other RCC histologies are frequently employed in tRCC without clear mechanistic rationale. Accordingly, response rates to these agents in tRCC are modest, highlighting the distinct biological features of this RCC subtype^7–10^. Indeed, genomic studies have revealed few recurrent genetic alterations in tRCC apart from the defining MiT/TFE fusion^3,11–13^. Most notably, tRCCs lack alterations in the *VHL* tumor suppressor gene; this contrasts with clear cell RCC (ccRCC), the most common type of RCC, in which loss of *VHL* and concomitant activation of hypoxia inducible factor 2 alpha (HIF-2a) are pathognomonic^3^. RNA-Seq studies have also suggested that tRCCs have a unique transcriptional profile and cluster distinctly from other RCCs in tumor datasets^3,14–16^.

Although is clear that the TFE3 fusion is the defining alteration in tRCC, the specific mechanism(s) by which it drives oncogenesis remain obscure. This is in contrast to driver fusions in other cancers (e.g. EWS-FLI, NRG1, RET), which have often been clearly linked to cancer hallmarks such as activating proliferation, growth factor signaling or invasion/metastasis^17–19^. While preclinical studies have nominated a few molecular pathways that may be activated in tRCC ^20–25^, these have not always been linked directly to the driver fusion. Indeed, a major barrier to developing effective therapies in tRCC has been an incomplete understanding of the oncogenic pathways driven by the TFE3 fusion.

RCCs are intrinsically metabolic diseases and several subtypes of kidney cancer are associated with distinctive metabolic dysregulations stemming from their underlying driver alterations ^26^. For example, activation of HIF signaling downstream of *VHL* loss in ccRCC results in metabolic reprograming with increased glycolysis and suppression of entry into the TCA cycle ^27,28^. In fumarate hydratase deficient RCC (FH-RCC), glycolytic metabolism is due to an intrinsic deficiency in the TCA cycle enzyme, fumarate hydratase (FH) ^29^. Birt Hogg Dube syndrome-associated chromophobe RCCs, which harbor folliculin (*FLCN*) inactivation, also shift metabolism toward aerobic glycolysis, but display increased mitochondrial mass secondary to peroxisome proliferator-activated receptor gamma coactivator 1-alpha (PGC1α) activation ^26^. Renal oncocytomas display inactivating mutations in mitochondrial complex I genes, leading to impaired oxidative phosphorylation (OXPHOS) ^30,31^. Alterations in succinate dehydrogenase (SDH), tuberous sclerosis (TSC) and fructose-1,6-bisphosphatase 1 (FBP1) represent additional lesions that drive metabolic reprogramming in kidney cancer^32^.

As these examples highlight, glycolytic metabolism is enforced by somatic mutation in most kidney cancer subtypes. Indeed, most cancers in general utilize glycolysis even under oxygen-replete settings where OXPHOS could be possible; this phenomenon of metabolic reprogramming, known as the “Warburg effect,” is a cancer hallmark ^33–36^. While some cancers do display evidence of enhanced OXPHOS in some contexts, this has not typically been linked to a genetic driver ^37^, and there are few examples of cancers that are for which aerobic respiration is a defining feature of the cancer’s metabolism. This raises the question of whether there are genetically-defined malignancies that represent the converse of highly glycolytic cancers.

While direct inhibition of bioenergetic pathways often has a narrow therapeutic window ^38^, recent studies have also indicated extensive crosstalk between metabolic reprogramming in cancer and pathways involved in redox homeostasis, which may inform additional vulnerabilities of specific metabolic states ^39^. For example, it has been recently shown that, while activation of the antioxidant regulator NRF2 factor (nuclear factor erythroid 2-like 2, encoded by the *NFE2L2* gene) is advantageous in lung cancers with glycolytic metabolism, its activation a subset of lung cancer cells high in OXPHOS decreases fitness by inducing NADH reductive stress and pushing the NADH/NAD+ ratio beyond a tipping point ^40,41^. This suggests that an overly reducing environment can prove to be a vulnerability in specific metabolic contexts. We have previously shown that tRCCs highly express some NRF2 target genes but curiously lack the somatic alterations in this pathway that are found in other kidney cancers ^42^, suggesting that tRCCs might exhibit a distinct interplay between bioenergetic preferences and redox homeostasis.

To date, the defining metabolic features of tRCC and their associated vulnerabilities remain unknown. In this study, we sought to understand the key metabolic phenotypes in tRCC, their mechanistic link to the TFE3 driver fusion, and their functional consequences.

### tRCCs display activation of OXPHOS metabolism

We recently performed comparative transcriptomics between tRCC tumors and other types of kidney cancer to identify pathways selectively activated in tRCC; this analysis revealed an enrichment for gene sets related to OXPHOS in tRCC^42^. To further extend this finding, we first assessed an OXPHOS transcriptional signature in three different RCC datasets (two of RCC tumors^14,43^ and one of RCC patient-derived xenografts^44^). In all three cohorts, we observed that tRCC tumors displayed heightened OXPHOS signatures relative to ccRCCs. By contrast, ccRCCs displayed higher glycolysis signature scores, consistent with the reliance of ccRCCs on aerobic glycolysis (Warburg effect)^45^ (Fig. 1a).

**Fig. 1:**
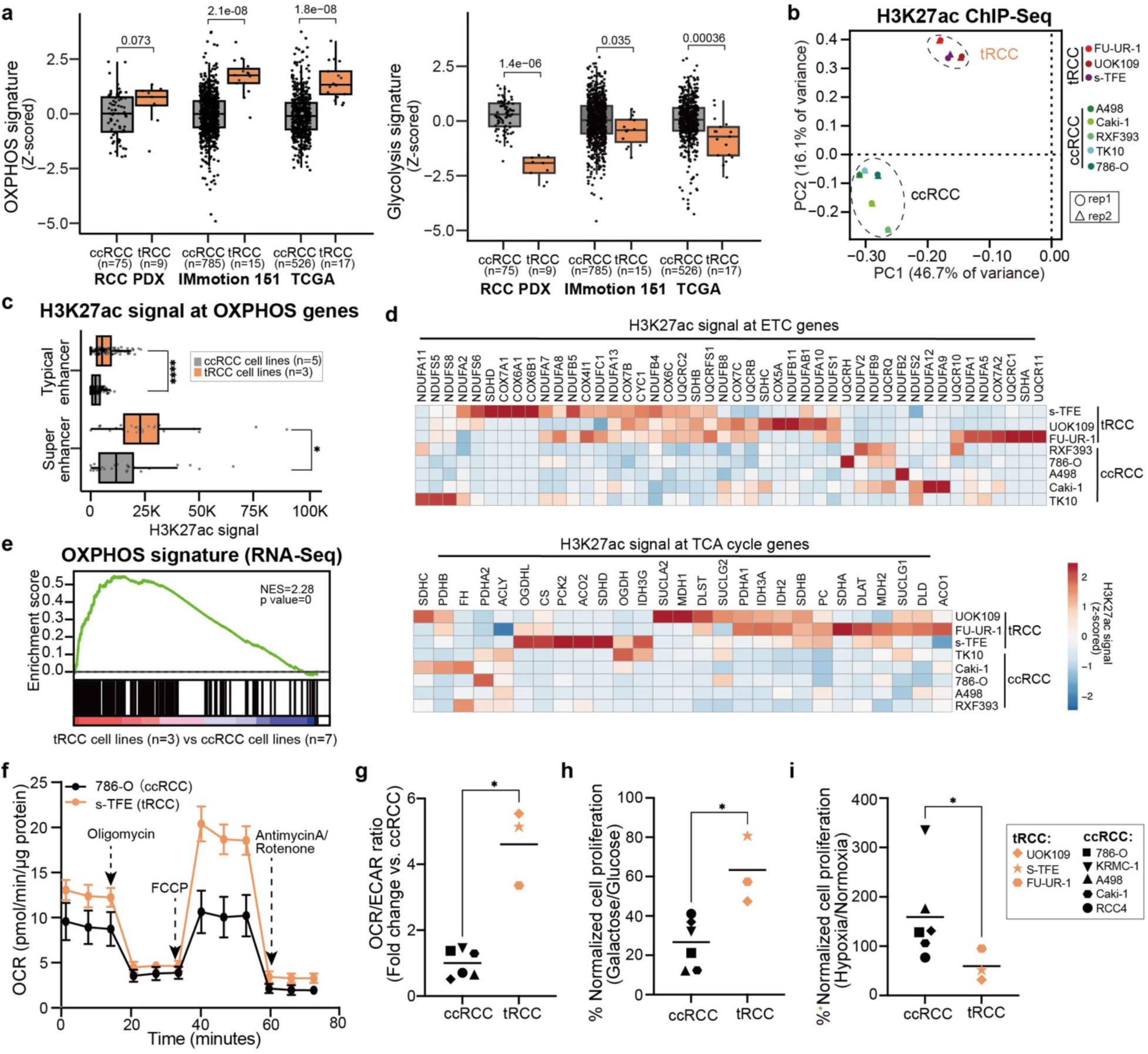
tRCCs bioenergetically favor OXPHOS. (a) OXPHOS and glycolysis gene signature scores in ccRCC or tRCC tumors from three independent studies (TCGA, Motzer et al., Elias et al. (PDX))^14,43,44^. (b) Principal component analysis (PCA) of H3K27ac ChIP-Seq data across 8 RCC cell lines (3 tRCC; 5 ccRCC; 4 ccRCC lines were profiled in a previously published study, see Extended Data Fig. S1^124^). (c) Boxplot of averaged H3K27ac signal at typical or super enhancers at OXPHOS genes in 3 tRCC cell lines (UOK109, FU-UR-1, s-TFE) vs. 5 ccRCC cell lines (786-O, Caki-1, RXF393, TK10, A498). (d) Heatmap showing H3K27ac signal (quantified by ROSE2) at ETC and TCA cycle genes in tRCC vs. ccRCC cell lines. (e) GSEA showing enrichment of OXPHOS gene signature in tRCC cell lines (n=3, UOK109, FU-UR-1, s-TFE) versus ccRCC cell lines from CCLE (n=7, A498, A704, 786-O, 769-P, Caki-1, Caki-2, OS-RC-2). (f) Oxygen consumption rate (OCR) as measured by a Seahorse Bioflux analyzer after the addition of oligomycin, FCCP, or antimycin A/rotenone in a ccRCC cell line (786-O) and a tRCC cell line (s-TFE). Data are shown as mean ± s.d, n=5 biological replicates for 786-O cell line, n=6 biological replicates for s-TFE cell line. (g) Ratio of (OCR) to extracellular acidification rate (ECAR) as detected by a Seahorse Bioflux analyzer in ccRCC (n=6, 786-O, Caki-1, Caki-2, KRMC-1, A498, RCC4) and tRCC (n=3, UOK109, FU-UR-1, s-TFE) cell lines. OCR/ECAR ratio represents the basal respiration:glycolytic balance in each cell line. Data are shown as mean ± s.d, n=5-7 biological replicates per cell line. (h) Viability of ccRCC (n=6, 786-O, Caki-1, Caki-2, KRMC-1, A498, RCC4) and tRCC (n=3, UOK109, FU-UR-1, s-TFE) cell lines cultured in glucose or galactose-containing media for 6 days. Data are shown as mean ± s.d. n=3 biological replicates per cell line. (i) Viability of ccRCC (n=6, 786-O, Caki-1, Caki-2, KRMC-1, A498, RCC4) and tRCC (n=3, UOK109, FU-UR-1, s-TFE) cell lines cultured under hypoxic (2.5% O_2_) or normoxic (20% O_2_) conditions for 10 days. Data are shown as mean ± s.d. n=3-4 biological replicates per cell line. For panels (a), (c) and (g-i), statistical significance was determined by Mann-Whitney U test. *p < 0.05, **p < 0.01, ***p < 0.001, **** p < 0.0001, n.s. not significant.

Multiple prior studies have indicated that differences in post-translational histone modifications can discriminate between cancer subtypes of a given lineage^46,47^. To identify distinguishing features of the epigenetic landscape in tRCC, we performed chromatin immunoprecipitation sequencing (ChIP-seq) for the active histone modification H3K27ac in 3 tRCC cell lines (UOK109, FU-UR-1, s-TFE) and 1 ccRCC cell line (786-O); we also integrated published H3K27ac ChIP-seq data for 4 additional ccRCC cell lines (A498, Caki-1, RXF393, TK10)^48^. Principal component analysis (PCA) of these 9 cell lines separated samples into two major groups. All ccRCC cell lines grouped together, while three tRCC cell lines (UOK109, FU-UR-1, s-TFE) formed a separate cluster (Fig. 1b and Extended Data Fig. 1a). This suggests that tRCCs have distinctive epigenetic features compared to ccRCC.

The H3K27ac modification marks active enhancers and is associated with active transcription. We used H3K27ac signal to annotate and rank active enhancers in each cell line across this panel (Table S1), distinguishing between typical enhancers (TEs) and superenhancers (SEs), the latter of which represent large enhancer clusters that activate the expression of oncogenic drivers^49,50^. We observed that H3K27ac signal at both TEs and SEs associated with OXPHOS pathway genes was higher in tRCC cell lines as compared with ccRCC cell lines (Fig. 1c). This included both genes in the tricarboxylic acid (TCA) cycle (the second stage in respiration responsible for the oxidation of acetyl-CoA and production of the reducing agents NADH and FADH2) as well as genes encoding multiple components of the electron transport chain (ETC), which is responsible for the oxidation of reduced electron carriers (NADH/FADH2) and transfer of electrons to oxygen via OXPHOS, leading to the generation of ATP (Fig. 1d and Extended Data Fig. 1b). Consistent with this epigenetic profiling data, RNA-Seq profiles of tRCC cells were enriched for an OXPHOS gene signature as compared with ccRCC cells (Fig. 1e). Together, these epigenomic profiling data suggest that aerobic respiration (OXPHOS) is transcriptionally driven in tRCC.

To determine the phenotypic correlates of these epigenomic features, we then investigated the bioenergetic preferences of tRCC cells. We determined the ratio of oxygen consumption rate (OCR) to extracellular acidification rate (ECAR) using a Seahorse metabolic flux analyzer. The OCR/ECAR ratio reflects a cell’s preference for mitochondrial respiration versus glycolysis, with a higher OCR/ECAR indicating a preference for OXPHOS (mitochondrial respiration)^51^. As an example, the tRCC cell line (s-TFE) displayed a distinct bioenergetic profile with markedly increased mitochondrial capacity compared to the ccRCC cell line, 786-O (Fig. 1f). Indeed, tRCC cells overall exhibited a significantly elevated OCR/ECAR ratio compared to ccRCC cells (Fig. 1g), indicating a relative preference for aerobic respiration in tRCC^41,52–54^. Consistent with this observation, tRCC cells were also more tolerant to growth in galactose-containing media, which forces cells to rely more heavily on OXPHOS for energy production^53,55,56^ (Fig. 1h).

The preference of tRCC cells to utilize aerobic respiration next prompted us to investigate whether this renders them more sensitive to growth under hypoxic conditions than ccRCC cells. Under conditions of hypoxia, OXPHOS is downregulated while glycolysis is activated^57,58^. ccRCC cells are adapted to hypoxic growth due to loss of the *VHL* tumor suppressor gene and resultant activation of the HIF-2α pathway, which renders ccRCCs highly glycolytic; however, *VHL* loss is not found in tRCC^42,59^. Indeed, we found that tRCC cells displayed impairment of growth under hypoxic conditions, while ccRCC cells were unaffected or increased proliferation under hypoxic conditions (Fig. 1i and Extended Data Fig. 1c). Together, these results indicate that, in contrast to ccRCCs that are highly glycolytic, tRCC cells rely prominently on aerobic respiration.

### Aerobic respiration in tRCC is transcriptionally driven by the TFE3 fusion

Since the TFE3 fusion is the defining (and often sole) genetic alteration in tRCC, we next sought to determine the link between the fusion and the metabolic features of tRCC. We performed Chromatin Immunoprecipitation and Sequencing (ChIP-Seq) using an antibody against the TFE3 fusion in three tRCC cell lines (FU-UR-1, ASPL-TFE3 fusion; s-TFE, ASPL-TFE3 fusion; UOK109, NONO-TFE3 fusion) and called high-confidence genomic binding sites (Methods and Table S2). We identified 1,347 TFE3 fusion peaks shared across all three tRCC cell lines (Fig. 2a).

**Fig. 2:**
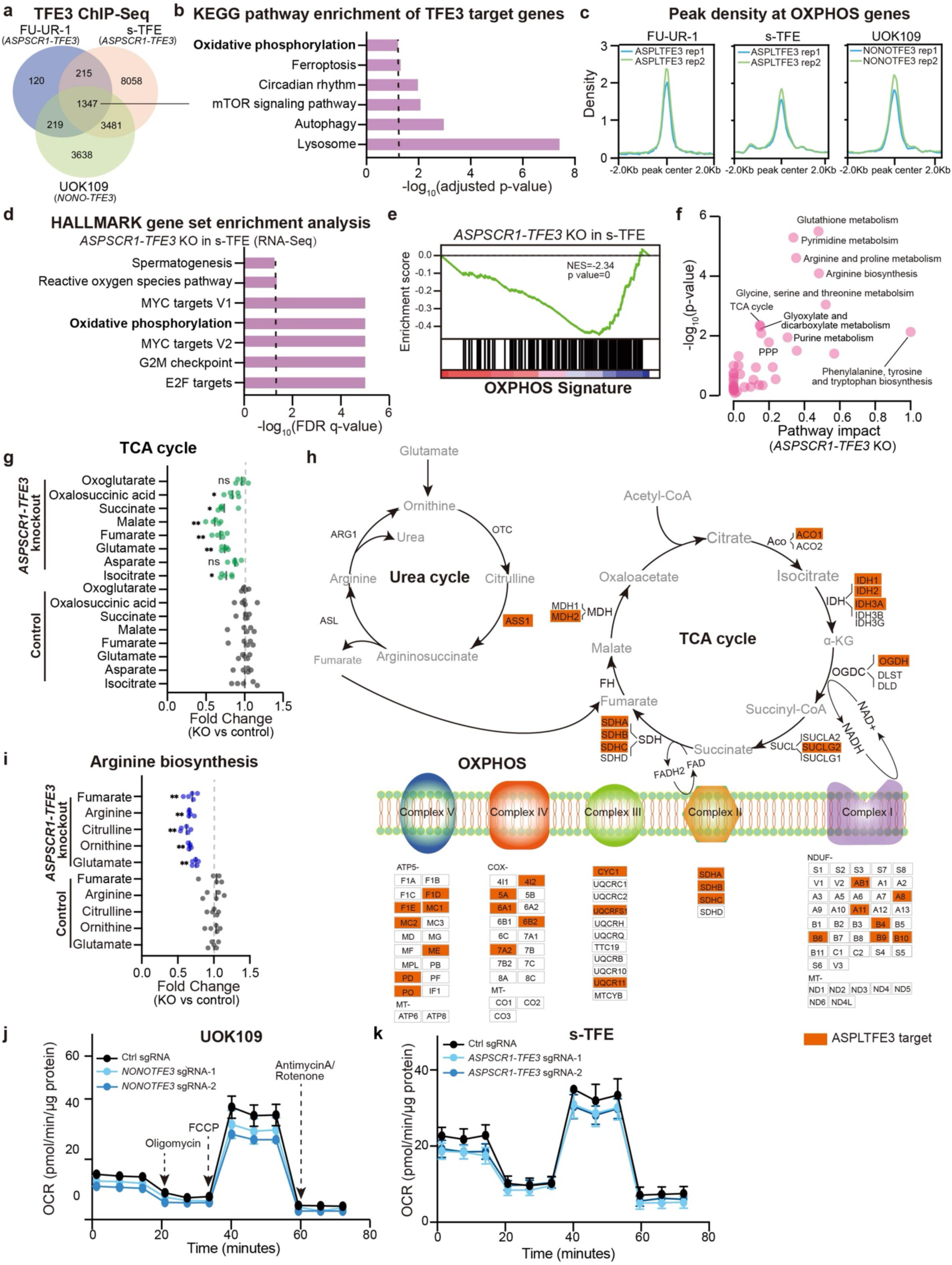
OXPHOS metabolism in tRCC is driven by the TFE3 fusion. (a) Venn diagram showing overlap of TFE3 fusion peaks detected by ChIP-Seq across three tRCC cell lines (FU-UR-1, s-TFE, UOK109). (b) KEGG pathway analysis showing pathways significantly enriched amongst genes proximal to TFE3 fusion peaks (from shared peaks in panel (a)). (c) Profile plot showing TFE3 fusion ChIP-Seq signal at OXPHOS genes. (d) Bar plot showing the top gene sets depleted upon *ASPSCR1-TFE3* knockout in s-TFE cells. (e) GSEA plot showing depletion of OXPHOS gene signature in s-TFE cells upon *ASPSCR1-TFE3* knockout. (f) KEGG analysis on untargeted metabolomic profiling data displaying metabolic pathways downregulated following *ASPSCR1-TFE3* knockout in s-TFE cells. (g) Change in levels of TCA cycle-related metabolites following *ASPSCR1-TFE3* knockout in s-TFE cells. For each metabolite, fold change was normalized to control sgRNA condition. Data are shown as mean ± s.d, n=5 biological replicates per cell line. (h) Schematic of urea cycle, TCA cycle, and the mitochondrial electron transport chain (ETC), annotated with genes that are ASPL-TFE3 targets as determined by ChIP-Seq in s-TFE cells (orange box). In the schematic, enzymes are in black text, metabolites are in gray text. (i) Change in levels of arginine biosynthesis-related metabolites following *ASPSCR1-TFE3* knockout in s-TFE cells. For each metabolite, fold change was normalized to control sgRNA condition. Data are shown as mean ± s.d, n=5 biological replicates per cell line. (j) Oxygen consumption rate (OCR) level after knockout of *NONO-TFE3* in UOK109 tRCC cell line. Data are shown as mean ± s.d, n=5-6 biological replicates. (k) Oxygen consumption rate (OCR) level after knockout of *ASPSCR1-TFE3* in s-TFE tRCC cell line. Data are shown as mean ± s.d, n=8-11 biological replicates. For panels (g) and (i), statistical significance was determined by Mann-Whitney U test. *p < 0.05, **p < 0.01, ***p < 0.001, **** p < 0.0001, n.s. not significant.

We then annotated the genes proximal to consensus TFE3 fusion peaks and subjected these genes to enrichment analysis. Top enriched pathways included lysosomal biogenesis, autophagy, and mTOR signaling, consistent with canonical features of wild type TFE3^60–62^. Notably, however, TFE3 fusion targets were also enriched for genes in OXPHOS metabolism (Fig. 2b). We examined enrichment of TFE3 fusion binding sites in proximity to OXPHOS-related genes and found strong TFE3 fusion binding in all three tRCC cell lines (Fig. 2c and Extended Data Fig. 2a). To determine whether expression of OXPHOS genes is regulated downstream of TFE3 fusion binding, we performed RNA sequencing (RNA-Seq) following *ASPSCR1-TFE3* knockout in s-TFE and FU-UR-1 tRCC cells. OXPHOS was among the most significantly downregulated pathways (Fig. 2d-e and Extended Data Fig. 2b-c). Protein levels of the mitochondrial respiratory complexes (e.g. complex I protein NDUFB8 and complex II protein SDHB) also decreased upon fusion knockout (Extended Data Fig. 2d). Overall, these results indicate transcriptional regulation of OXPHOS-related genes by the TFE3 fusion through direct binding at genes critical for aerobic respiration (Fig. 2b).

To further characterize the metabolic profile driven by the fusion, we performed untargeted metabolic profiling after *ASPSCR1-TFE3* knockout in s-TFE cells (Methods) and subjected differentially abundant metabolites to pathway analysis. Among the pathways most significantly impacted by fusion knockout, we noted the TCA cycle and arginine biosynthesis/metabolism (Fig. 2f). Analysis of metabolite levels within both the TCA cycle and the urea cycle (responsible for arginine synthesis as well as fumarate production, the latter of which enters the TCA cycle)^63–65^ revealed a significant downregulation of metabolites in both pathways upon *TFE3* fusion knockout (Fig. 2g and 2i). Integration with our ChIP-Seq data revealed that several critical genes in the TCA cycle, urea cycle, and ETC are direct transcriptional targets of the TFE3 fusion (Fig. 2h, orange boxes). We also noted that the expression of the urea cycle enzyme arginosuccinate synthetase 1 (*ASS1*), which lays upstream of fumarate production and is typically suppressed in ccRCC^66^, is activated by TFE3 fusions and highly expressed in tRCC (Extended Data Fig. 2e). Finally, we performed Seahorse metabolic flux analysis and observed a decrease in OCR upon *TFE3* fusion knockout in tRCC cells (Fig. 2j-k and Extended Data Fig. 2f).

Interestingly, knockout of wild type *TFE3* in a ccRCC cell line (786-O) did not lead to decreased expression of OXPHOS related genes (Extended Data Fig. 2g), nor were there effects on TCA cycle or arginine biosynthesis-related metabolites (Extended Data Fig. 2h-i) or on OCR (Extended Data Fig. 2j), suggesting that this activity may be a selective property of the constitutively active TFE3 fusion. Together, these results indicate metabolic reprogramming toward aerobic respiration is under direct transcriptional control of the driver TFE3 fusion in tRCC.

### Metabolic reprogramming by TFE3 fusions renders tRCC cells sensitive to reductive stress

Almost all subtypes of RCC maintain high glutathione levels, partially reflective of their origin in kidney tubular cells ^67^, and the mechanisms by which these high GSH levels are maintained differ between subtypes ^30,68,69^. Baseline metabolite profiling revealed that tRCCs, like ccRCCs, have a high ratio of reduced to oxidized glutathione (GSH/GSSG) (Extended Data Fig. 3a). Glutathione metabolism was the top pathway downregulated upon *ASPSCR1-TFE3* knockdown in s-TFE cells upon metabolite profiling; we also observed downregulation of the PPP, which generates the reducing equivalent NADPH and plays a role in maintaining redox balance (Fig. 2f). To investigate whether these pathways might be directly regulated by the TFE3 fusion in tRCC, we examined fusion binding sites proximal to genes involved in glutathione metabolism or the PPP. We observed strong fusion binding proximal to these genes, with key enzymes in both pathways being direct TFE3 fusion transcriptional targets (Extended Data Fig. 3b-c). Moreover, most of these genes were transcriptionally downregulated on RNA-Seq after *ASPSCR1-TFE3* knockout (Extended Data Fig. 3d), as were key metabolites in these pathways (Extended Data Fig. 3e). Finally, baseline ROS levels were significantly lower in tRCC cells than in ccRCC cells (Extended Data Fig. 3f) and increased upon knockout of the fusion (Extended Data Fig. 3g).

In other RCCs, which are glycolytic, somatic activation of the NRF2 pathway can drive flux through pathways that produce reducing equivalents; however, such somatic alterations are not found in tRCC^3,70^. While we did observe expression of nuclear NRF2 in tRCC cells (Extended Data Fig. 3h-i), NRF2 levels were lower than typically seen with NRF2 mutation or *KEAP1* inactivation. In many cases, NRF2 appeared activated secondary to high levels of *SQSTM1* (which encodes the p62 autophagy receptor protein, a TFE3 fusion target) (Extended Data Fig. 3j-m). Moreover, unlike highly glycolytic cells with high NRF2 activity, which are uniquely sensitive to NRF2 inhibition^41,71^, tRCC cells are only modestly sensitive to *NFE2L2* knockdown (^42^ and data not shown). Altogether, these results indicate that the TFE3 fusion drives a highly reductive environment by directly activating transcription of multiple genes involved in the production of the reducing equivalents glutathione, NADH, and NADPH.

While an increased GSH/GSSG ratio and elevated levels of reduced nucleotide cofactors (e.g. NADH and NADPH) can help to detoxify ROS, heightened levels of antioxidants can also result in an overly reductive environment that can be detrimental to cell fitness by priming a cell to “reductive stress”^40^. Reductive stress can encompass multiple flavors, all resulting from an imbalance of metabolic reducing equivalents, including NADH-reductive stress (elevated NADH/NAD+) and glutathione (GSH)-reductive stress (elevated GSH/GSSG)^40,72^. In the context of lung cancer, it has been recently shown that cells with heightened OXPHOS and low glycolytic metabolism are particularly reliant on Complex I of the ETC for NADH oxidation, and are therefore vulnerable to NADH reductive stress caused by NRF2 pathway activation^41^.

We therefore sought to determine whether tRCC cells – which we have shown to have enhanced OXPHOS and a highly reducing environment directly driven by the TFE3 fusion – represent a cancer type vulnerable to reductive stress. Indeed, we observed that overexpression of NRF2 in tRCC cells impaired proliferation, although it enhanced the proliferation of highly glycolytic ccRCC cells (Extended Data Fig. 4a-b). tRCC cells were also more sensitive to the knockout of *KEAP1*, which activates NRF2 signaling, than were ccRCC cells (Extended Data Fig. 4c-d). It has been previously shown that NADH levels are both necessary and sufficient for NRF2 sensitivity^41^. In agreement with this prior study^41^, we observed that NRF2 overexpression led to an increased NADH/NAD+ ratio (as measured by a genetically-encoded NADH/NAD+ reporter, SoNar)^73^ in tRCC cells (FU-UR-1 and s-TFE) compared to ccRCC cells (786-O) (Extended Data Fig. 4e-f). Overexpression of the NADH-oxidizing enzyme LbNOX^74^ partially rescued the detrimental effects of NRF2 induction on proliferation of tRCC cells (Extended Data Fig. 4g-h). Finally, knockout of the TFE3 fusion led to an increased NADH/NAD+ ratio by SoNar (Extended Data Fig. 4i), presumably mediated through the multiple OXPHOS genes that are TFE3 transcriptional targets (Fig. 2h). Altogether, our results indicate that TFE3 fusions drive a transcriptional program that promotes a highly reductive cellular environment. While this antioxidant program may be beneficial in counterbalancing the ROS generated by OXPHOS, it also creates a unique metabolic state that renders tRCCs vulnerable to reductive stress.

### TFE3-dependent metabolic reprogramming evinces EGLN1 as a druggable dependency in tRCC

We then leveraged genome-scale CRISPR knockout screening to systematically uncover metabolic dependencies in s-TFE tRCC cells. We quantified gene dependencies in this cell line as previously described^75^ and compared dependency scores by gene to five well-annotated ccRCC cell lines subjected to CRISPR screening in a published effort^76,77^. We found that s-TFE cells displayed higher gene dependency scores across 198 OXPHOS-related genes as compared with ccRCC cells (Fig. 3a). Direct inhibition of OXPHOS via Complex I inhibition has been tested in humans and has a narrow therapeutic window with serious dose-limiting toxicities ^38,78^. Therefore, we sought to interrogate our CRISPR screening data to identify additional selective metabolic vulnerabilities in tRCC that might converge on this pathway. We overlapped the top 1000 gene dependencies in s-TFE or ccRCC cell lines (averaged across 5 ccRCC cell lines screened in the Cancer Dependency Map) with lists of druggable genes^71^ and metabolic genes^79^. This resulted in a set of 48 druggable metabolic dependencies in s-TFE cells and 39 in ccRCC cells. We compared dependency scores of these genes in s-TFE vs. ccRCC cells; this revealed *EGLN1* as the strongest selective dependency of tRCC cells, with no dependency in ccRCC cells (Fig. 3b). *EGLN1* encodes a prolyl hydroxylase enzyme that acts as a critical oxygen sensor by hydroxylating the α subunit of the hypoxia-inducible factor (HIF-1α) and targeting it for degradation by the VHL complex^80,81^.

**Fig. 3:**
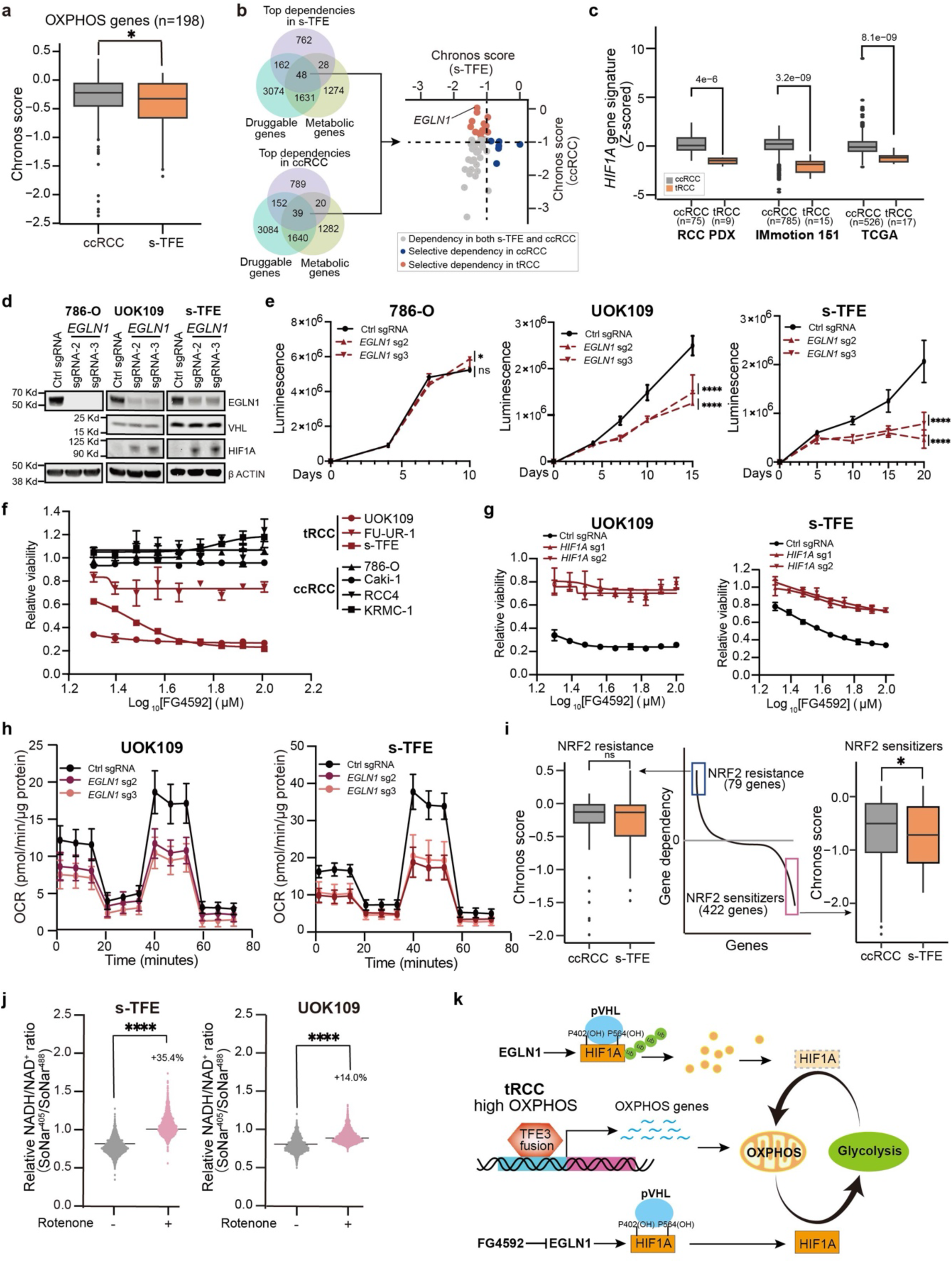
Selective *EGLN1* dependency in tRCC. (a) Comparison of dependency scores for OXPHOS genes in ccRCC cell lines (averaged across 5 cell lines for each gene) vs. s-TFE tRCC cells. (b) Gene dependencies identified via genome-scale CRISPR/Cas9 screening in s-TFE (tRCC) or ccRCC cells were overlapped with druggable and metabolic gene lists to nominate *EGLN1* as a candidate selective dependency in tRCC cells. (c) *HIF1A* gene signature scores in ccRCC or tRCC tumors from three independent studies as in Figure 1A. (d) Western blot showing expression of EGLN1, VHL, and HIF1A after knockout of *EGLN1* in a ccRCC cell line (786-O) or tRCC cell lines (UOK109 and s-TFE). (e) Cell proliferation of tRCC cell lines (UOK109 and s-TFE) and ccRCC (786-O) after knockout of *EGLN1*. Data are shown as mean ± s.d. n=3 biological replicates. (f) Relative cell viability (normalized to DMSO control) of tRCC cell lines or ccRCC lines after treatment with the EGLN inhibitor FG4592. Data are shown as mean ± s.d. n=3 biological replicates. (g) Relative viability of *HIF1A* knockout tRCC cell lines (n=2, UOK109 and s-TFE) after treatment with FG4592. Data are shown as mean ± s.d. n=3 biological replicates. (h) OCR after knockout of *EGLN1* in UOK109 and s-TFE cell lines. Data shown as mean ± s.d, n=8-16 biological replicates. (i) Genes whose knockout has been previously reported^41^ to confer sensitivity (422 genes) or resistance (79 genes) to NRF2 activation were compared for dependency in s-TFE cells vs. ccRCC cells (average of dependency score for 5 ccRCC cell lines, for each gene). (j) Quantification of NADH to NAD+ ratio via SoNar assay following rotenone treatment in tRCC cell lines (UOK109 and s-TFE). (k) Model: EGLN1 modulates OXPHOS and is a dependency in tRCC. For panel (a), (c), (e), and (i-j), statistical significance was determined by Mann-Whitney U test. *p < 0.05, **p < 0.01, ***p < 0.001, **** p < 0.0001, n.s. not significant.

We hypothesized that *EGLN1* knockout would lead to HIF-1α stabilization selectively in tRCC cells, given that *VHL* is retained in this RCC subtype and because 14q deletion (where the *HIF1A* gene is located) is uncommon in tRCC, in contrast to ccRCC^42,82^. Indeed, we observed that, despite higher levels of *HIF1A* mRNA in tRCC vs. ccRCC tumors, a HIF1A gene signature was relatively suppressed in tRCC tumors, consistent with active degradation of HIF-1a by the EGLN1-VHL axis in tRCC (Fig. 3c and Extended Data Fig. 5a-b). As HIF-1α functions a metabolic switch to reprogram metabolism away from OXPHOS and toward glycolysis, we reasoned that *EGLN1* knockout could stabilize HIF-1α and thereby have a detrimental effect on tRCC cells, which we have shown to be uniquely dependent on OXPHOS^83,84^.

Indeed, we observed that knockout of *EGLN1* led to stabilization of HIF-1α protein selectively in tRCC cells (Fig. 3d) (VHL and HIF1a were not detectable in 786-O cells). This was accompanied by a selective impairment of cell proliferation with *EGLN1* knockout in tRCC cells but not ccRCC cells (Fig. 3e). Consistently, knockout of *VHL*, which would prevent degradation of EGLN1*-*hydroxylated HIF-1α, was also selectively essential in tRCC cells expressing high *HIF1A* (Extended Data Fig. 5c-f); *VHL* was dispensable both in the *VHL*-null ccRCC line 786-O and in the *VHL-* preserved ccRCC line Caki-1^77^ (Extended Data Fig. 5g-h). This effect was phenocopied by Roxadustat (FG-4592), an orally-available EGLN inhibitor and HIF stabilizer^85,86^, and almost completely rescued by concurrent *HIF1A* deletion (Fig. 3f-g and Extended Data Fig. 5i). Consistent with the role of HIF-1α as a metabolic switch between OXPHOS and glycolysis, knockout of both *EGLN1* and *VHL* markedly reduced OCR in tRCC cells (Fig. 3h and Extended Data Fig. 5j). We conclude that *EGLN1* represents a strong and selective vulnerability in tRCC, likely driven by a mechanism of HIF-1α stabilization that results in metabolic reprogramming away from OXPHOS and toward glycolysis, which is detrimental to tRCC cells (Fig. 3k).

To gain further insight into the mechanism by which OXPHOS inhibition (either by TFE3 fusion inhibition or EGLN1 inhibition) could impair fitness in tRCC, we overlapped gene dependencies in s-TFE cells with hits from a prior CRISPR screen that identified modulators of NRF2 sensitivity^41^. We found that gene dependencies in s-TFE cells showed strong overlap with genes whose knockout was found to sensitize to NRF2 hyperactivation in this prior study^41^. Notably, however, this was not the case in ccRCC cell lines (Fig. 3i). Treatment of tRCC cell lines with the mitochondrial Complex I inhibitor rotenone led to an increased NADH/NAD+ ratio by SoNar, as did knockout of *TFE3* fusions (Fig. 3j and Extended Data Fig. 4i). These results suggest that impinging on the OXPHOS pathway in tRCC induces reductive stress.

Recent genomic studies have implicated a distinct biology driving tRCC as compared with other RCC subtypes^3,11–13,87^. While the MiT/TFE gene fusion represents the pathognomonic genetic lesion in tRCC, the precise molecular mechanisms by which it drives cancer have remained obscure. Here, our integrative analysis of multiple cell line models and tRCC tumors converges on metabolic reprogramming towards OXPHOS as a shared and critical oncogenic function of TFE3 fusions. This work clearly places tRCC alongside other RCCs as being typified by mutation-driven metabolic dysregulation^26^. However, our results clarify that tRCC has a metabolic profile that is distinct from other kidney cancers. This is of clinical relevance, given that tRCCs are frequently histologically misclassified as either ccRCC or papillary RCC; our study suggests that therapies targeted to those subtypes may not necessarily be active in tRCC, owing to its unique biology ^7,8,10^.

We show that tRCCs bioenergetically favor OXPHOS, a program that is driven directly by the TFE3 fusion. While this is in stark contrast to most other RCCs, in which the TCA cycle is suppressed, a recent study has interestingly shown that ccRCCs may shift to a mitochondrial respiration program during metastasis^88^. Metastatic rates for RCC vary by subtype, with certain subtypes metastasizing from smaller primary tumors^89^. We speculate that the proclivity of tRCCs to aerobic respiration may be linked to their inherent aggressiveness and propensity to metastasize early^90^. Interestingly, a recent study reported an OXPHOS transcriptional program in alveolar soft part sarcoma (ASPS), a distinct cancer also driven by TFE3 fusions; this suggests that the TFE3 fusion may drive similar metabolic programs across cancer types^91^. Consistent with this notion, *Tfe3* knockout mice exhibit altered mitochondrial morphology and dynamics, with impaired mitochondrial respiratory function ^92^.

Most cancers show evidence of aerobic glycolysis, and in several kidney cancers, a deficiency in aerobic respiration appears to be selected for by genetic mutation (e.g. *FH* mutations in *FH-RCC* or Complex I mutations in renal oncocytoma). By contrast, tRCC appears to represent a unique case in which OXPHOS is induced by the driver fusion and is presumably under positive selection. This cancer may therefore represent an attractive model in which to study aerobic respiration driven by genetic alteration, and the mechanisms by which such an event might be oncogenic. More broadly, while rewiring of metabolism is a cancer hallmark, there are a limited number of examples in which these phenotypes are tightly linked to genetic biomarkers. The fact that an OXPHOS program appears directly attributable to the singular genetic event in tRCC (the TFE3 fusion) opens the possibility that this axis can be therapeutically exploited.

Recent studies have suggested that inhibiting OXPHOS in cancer cells may be a viable therapeutic strategy^37^. Pharmacologically, this can be achieved through selective mitochondrial complex I inhibitors^93^, or via the anti-diabetic drug metformin, which has activity on complex I^94,95^. Unfortunately, despite promising preclinical studies, a recent clinical trial suggested that IACS-010759, a potent and selective complex I inhibitor, had a narrow therapeutic window with serious dose-limiting toxicities, warranting caution for targeting OXPHOS via this mechanism in cancer^38,78^. Our results suggest that EGLN1 inhibition may be a means to more selectively shift the balance between OXPHOS and glycolysis in tRCC cells. Notably, *EGLN1* dependency is quite selective across cancers^96^ and multiple EGLN inhibitors have been clinically developed, approved, and are generally well-tolerated in humans^85,86,97,98^.

We show that, in addition to driving OXPHOS, TFE3 fusions activate a host of genes involved in glutathione and NADPH biosynthesis, thus establishing a highly reducing cellular environment that is linked to unique metabolic vulnerabilities. High levels of glutathione are generally associated with resistance to cell death via ferroptosis, but interestingly, perturbing OXPHOS or ETC function sensitizes to death via ferroptosis^99,100^. The highly reducing environment in tRCC, in conjunction with aerobic respiration driven by the TFE3 fusion, also makes this RCC subtype uniquely vulnerable to reductive stress. This metabolic liability can be exploited therapeutically by either inhibition of OXPHOS or hyperactivation of NRF2 signaling. While TFE3 fusions activate certain NRF2 target genes involved in the antioxidant response, it is notable that somatic alterations in the NRF2 pathway (e.g. *KEAP1* inactivation, *NFE2L2* activation, chr5q gain) are not found in tRCC, unlike other RCCs^3,101–103^. This suggests that somatic activation of NRF2 may not be tolerated in tRCC owing to their dependence on aerobic respiration, as such events would be predicted to constrain aerobic respiration, upset NADH/NAD+ balance, and result in NADH-reductive stress^41,104^.

Our work illuminates actionable metabolic features driven by the TFE3 fusion in tRCC that are distinct from other RCC subtypes, offering hope that molecularly-directed therapies can be advanced to specifically target the biology of this aggressive subtype of kidney cancer.

## Methods

### Cell culture

786-O (ATCC, CatLog: ATCC^®^ CRL-1932 ^TM^), 293T (ATCC, CatLog: ATCC^®^ CRL-11268^TM^), RCC4 (Sigma: #3112702), KRMC-1(JCRB, CatLog:JCRB1010), A498(ATCC, CatLog: ATCC^®^ HTB-44^TM^), Caki-1(ATCC, CatLog: ATCC^®^ HTB-46^TM^), Caki-2 (ATCC, CatLog: ATCC^®^ HTB-47^TM^), UOK109 (Dr. W. Marston Linehan’s laboratory, National Cancer Institute), FU-UR-1(Dr. Masako Ishiguro’s laboratory Fukuoka University School of Medicine) and s-TFE(RIKEN, # RCB4699 were grown at 37°C in DMEM supplemented with 10% FBS, 100 U mL^−1^ penicillin, and 100 μg mL^−1^ Normocin (Thermo fisher: #NC9390718).

### Antibodies

H3K27ac (Diagenode, Cat#: C15410196; RRID: AB_2637079), NRF2 (Cell Signaling Tech, Cat#: 12721; RRID: AB_2715528), NRF2 (Abcam, Cat#: ab62352; RRID: AB_944418), TFE3 (Sigma, Cat#: HPA023881; RRID: AB_1857931), TFE3 (Sigma, Cat#: SAB4200824; RRID: N/A), p62 (Cell Signaling Tech, Cat#: 5114S; RRID: AB_10624872), HIF1A (Fisher Scientific, Cat#: BDB610959; RRID: N/A), EGLN1 (Cell Signaling Tech, Cat#: 4835S; RRID: AB_10561316), VHL (Cell Signaling Tech, Cat#: 68547; RRID: AB_2716279), ACTIN (Cell Signaling Tech, Cat#: 8457; RRID: AB_10950489), ACTIN (Cell Signaling Tech, Cat#: 3700; RRID: AB_2242334), V5 (Life Technologies, Cat#: R96025; RRID: AB_2556564), Mouse IgG (Santa Cruz, Cat#: sc-2025; RRID: AB_737182), Rabbit IgG (Cell Signaling Tech, Cat#: 2729; RRID: AB_1031062), Donkey anti-Rabbit IgG (H+L) (Life Technologies, Cat#: A10042; RRID: AB_2534017), kEAP1 (Cell Signaling Tech, Cat#: 4678S; RRID: AB_10548196), OXPHOS (Abcam, Cat#: ab110411; RRID: AB_2756818)

### Plasmid construction

All ORFs were cloned into the pLX403(Addgene, #41395 puromycin resistance; #158560, blasticidin resistance) by Gibson cloning or Gateway cloning. All doxycycline inducible shRNA were cloned into a Gateway-compatible lentivector pLV706^105^. All sgRNAs were cloned into plentiCRISPRv2 (Addgene, #52961, puromycin resistance). Target sequences for shRNAs and sgRNAs are listed in Table S4. All the constructs were confirmed by Sanger sequencing.

### RNA extraction and RT-qPCR

ccRCC cell line (786-O) and tRCC cell line (FU-UR-1) were transduced with lentivirus expressing doxycycline shRNA against wild type TFE3 and ASPSCR1-TFE3 and selected with 500 mg/mL of G418. Subsequently, the cells were treated with doxycycline at a concentration of 1mg/mL for 5 days. s-TFE tRCC cells were transduced with control sgRNA or sgRNA targeting ASPSCR1-TFE3 for 5 days. Total RNA was isolated using RNeasy Plus Mini Kit (QIAGEN, #74136).

### Western blot analysis

Cells were resuspended in RIPA lysis buffer (Thermo Fisher Scientific, #89901) supplemented with protease inhibitors (Roche, #11836170001) and phosphatase inhibitors (Roche, #4906845001) on ice for 30 min. Total soluble protein was obtained by centrifugation at 12,000 rpm at 4°C for 15 min. The concentration of protein was measured using Pierce BCA protein Assay Kit (Thermo Fisher Scientific, #23225). Equal amounts of protein were loaded onto NuPAGE 4-12% Bis-Tris Protein Gels (Thermo Fisher Scientific, #NP0335) for separation by SDS-PAGE. Proteins were transferred to nitrocellulose membranes (Life Technologies, #IB23001) using an iBlot2 (Thermo Fisher Scientific), The nitrocellulose membranes were blocked with blocking buffer (Thermo Fisher Scientific, # NC1660550). Immunoblot analysis was performed with the indicated primary antibodies in antibody dilution buffer (Thermo Fisher Scientific, #NC1703226) overnight at 4°C. Membranes were incubated with secondary antibodies in antibody dilution buffer. Membranes were imaged using the Odyssey Clx Infrared Imaging System (LI-COR Biosciences).

### Chromatin immunoprecipitation (ChIP)

ChIP was performed as previously described^106^. Briefly, 3 x 10^6^ cells were fixed using 1% formaldehyde (Life technology: #28906) without methanol for 5 minutes at room temperature followed by quenching with 125 mM glycine (Sigma-Aldrich: #50046). Cells were washed twice with PBS and resuspended in 130 μL SDS lysis buffer. Chromatin was sheared to 200-400 bp using a Covaris E220 sonicator and cleared by centrifugation for 15 min at 13,000 rpm. 100 μL sample was diluted ten-fold with ChIP dilution buffer, and then incubated with protein A and protein G dynabeads (1:1 mix) and indicated antibody (H3K27ac, Diagenode, #C15410196; TFE3, Sigma, #HPA023881) at 4c overnight. Antibody-bound DNA was subsequently washed with 1 mL low salt buffer, 1 mL high salt buffer, 1 mL LiCl buffer once, respectively, and then washed twice with TE wash buffer. ChIP DNA was reverse-crosslinked and purified for DNA library construction. ChIP DNA library was made using NEBNext^®^ Ultra™ II DNA Library Prep Kit (New England Biolabs, #E7645S).

### Seahorse assay

Oxygen consumption rates (OCR) and extracellular acidification rates (ECAR) were determined with the XF Cell Mito Stress Kit (Agilent: #103015-100). All tRCC cell lines were seeded on a poly-L-lysine coated 96-well Seahorse plate (Agilent: #101085-004) at 4.5 x 10^3^ cells/well. ccRCC cell lines were seeded at 3.5 x 10^3^ cells/well (786-O, Caki-1, Caki-2, KRMC-1, A498) and 3 x10^3^ cells/well (RCC4). Cells were incubated overnight at 37 c in 5% CO2 incubator. XF96 FluxPak sensor cartridge was hydrated according to manufacturer’s instructions. During the following days, the growth medium was removed, and cells were washed with pre-warmed seahorse medium (XF DMEM (Agilent: #103575-100) supplemented with 10 mM glucose, 1 mM pyruvate solution, and 2 mM glutamine). After washing, cells were incubated in 180 μL Seahorse medium at 37 c in non-CO2 incubator for 45-60 min. The oxygen consumption rates were measured by XFe 96 extracellular flux analyzer by adding oligomycin, FCCP and rotenone/antimycin A to each cartridge port. All OCR values were normalized to total protein content as measured by BCA (Thermo Fisher Scientific, #23225) according to manufacturer’s instructions.

### Confocal microscopy

ccRCC cell line (786-O) and tRCC cell lines (UOK109, FU-UR-1, s-TFE) cells were cultured on glass coverslips at 37 c in 5% CO2 incubator for 24h. Cells were rinsed twice with pre-cold PBS and fixed with 4% paraformaldehyde for 10 min. 0.5% Triton X-100 was added to cells for 10 min for permeabilization. Subsequently, coverslips were blocked in 2.5% BSA in PBS for 1h and incubated with primary antibody (diluted in 2.5% BSA) at 4°C overnight. Following overnight incubation, coverslips were washed three times with TBS-T, then incubated with secondary antibody at room temperature for 1h. Coverslips were then mounted with DAPI medium and images were obtained with a 63x oil objective.

### Live cell imaging, image segmentation and quantifications

For NRF2 overexpression experiments, ccRCC cell line (786-O) and tRCC cell lines (FU-UR-1, s-TFE) expressing SoNar reporter and dox-inducible NRF2 were seeded on a glass bottom 96 well plate (iBidi) at 7 x 10^3^ cells/well for 1 day and then treated with dox the following day for 9h. For rotenone treatment experiments, tRCC cell lines (UOK109 and s-TFE) were seeded on a glass bottom 96 well plate (iBidi) at 7 x 10^3^ cells/well for 1 day and then treated with rotenone for 30 minutes. For *TFE3* fusion or *EGLN1* knockout experiments, the indicated tRCC cell lines expressing control sgRNA, *TFE3* fusion sgRNA or *EGLN1* sgRNA were seeded on a glass bottom 96 well plate (iBidi) at 7 x 10^3^ cells/well for 1 day. Subsequently, all plates were aintained at 37 c in CO2 incubator and imaged with the Leica Thunder 3D Cell Culture imager using the HC PL APO 40x/1.10 water objective and a Hamamatsu ORCA-Flash4.0 LT3 Digital CMOS camera. To assess relative NAD+ level, cells expressing SoNar were excited with a 488 nm light and emission was recorded within the 500–570 nm wavelength range. For relative NADH levels, a 395 nm light was used for excitation and emission was measured from a 500–570 nm wavelength range. The images were then processed using CellProfiler to calculate NADH/NAD+ ratio. For image segmentation and quantifications, cells were segmented using Cellpose^107^ plugin in CellProfiler^108^. Segmentation masks were generated using Cellpose pre-trained model “cyto2” and a cell diameter of 350 pixels (minimum size of 250 pixels and flow threshold 0.7). The segmented masks and original images were then fed into CellProfiler to quantify mean intensity value for both NADH and NAD+. Ratiometric images of SoNar were subjected to processing in Image J, wherein they were converted into 32-bit images and subsequently rendered in a 16-color mode for presentation.

### Intracellular ROS measurement

ccRCC and tRCC cells were seeded in a 6-well plate for 24h. Cells were washed twice with PBS, and then incubated with pre-warmed PBS buffer containing 5 μM CM-H2DCFDA (Thermo Fisher: #C6827) probe for 30 min. Subsequently cells were cultured in complete growth medium at 37 c incubator with 5% CO2 for 30 min. Cells were collected and resuspended in FACS buffer (PBS+2%FBS). The fluorescent signal at 530nm following excitation at 488nm was measured using a Fortessa flow cytometer.

### Cell proliferation assay

For CRISPR knockout experiments, the indicated cell lines were transduced with indicated sgRNA. Subsequently, cells were seeded in 96-well plates at varying densities ranging from 200-2000 cells per well, depending on the cell line. At the indicated time point, cell growth medium was removed from plates and the Cell Titer Glo reagent (Promega, #G7571) was added. The plates were then incubated on a shaker at room temperature for 10 minutes. The luminescence was measured on SpectraMax plate reader. For cell proliferation assays in glucose and galactose containing-DMEM culture medium, the indicated ccRCC and tRCC cells were seeded in a 96-well plate at varying densities ranging from 1000 to 3000 cells per well in glucose or galactose-containing medium, depending on the cell line. After 6 days, cells were fixed and stained with Hoechst, cell numbers were determined using a Celigo Imaging Cytometer.

### Cell growth assay under hypoxia or normoxia condition

The indicated cells were seeded in two 12 well plates separately, with densities ranging from 500 to 6000 cells per well depending on the cell lines. The normoxia plate was put into at 37 c incubator with 5% CO2 and 20% O2 for 10 days, while the hypoxia plate was put into 37 c incubator with 5% CO2 and 2.5% O2 for 10 days. Following the incubation period, the cells were fixed and stained with crystal violet. To quantify results, the crystal violet was destained with 10% acetic acid and absorbance was measured at 590 nm.

### Colony formation assay

*NFE2L2* and *LbNOX* were cloned to a doxycycline inducible pLX403 vector by gateway cloning. The indicated cell lines were transduced with lentivirus expressing *NFE2L2* or *LbNOX* and selected with 5 μg/mL of puromycin. Subsequently, the indicated cells were seeded in a 12 well plate, with densities ranging from 800 to 3000 cells per well depending on the cell line. Every two days, the culture medium was refreshed with or without the addition of 1μg/mL doxycycline. For *KEAP1* knockout experiments, the indicated cell lines were transduced with lentivirus expressing control sgRNA or *KEAP1* sgRNA selected with 5 μg/mL of puromycin, then the indicated cells were seeded in a 12 well plate, with densities ranging from 800 to 3000 cells per well depending on the cell line. After 12-25 days, the cells were fixed and stained with crystal violet. To quantify the results, the crystal violet was destained with 10% acetic acid and absorbance was measured at 590 nm.

### Drug treatment assay

The indicated cells were seeded in 96-well plates, ranging from 500 to 3000 cells per well depending on the cell line. EGLN1 inhibitor FG4592 (Selleckchem, #S1007) was added at the indicated concentration with a D330e Digital Dispenser (Tecan), with DMSO as a negative control. After 3 days, the medium was replaced with fresh medium and the EGLN1 inhibitor was re-added at the same concentration. After 7 days, cell viability was measured with the Cell Titer-Glo luminescent Cell Viability Assay (Promega, #G7571).

### Metabolic fingerprinting

ccRCC (786-O, A498, RCC4) and tRCC (FUUR-1, s-TFE, UOK109) cells were used. For knockout experiments, cells were transduced with pLenti-CRISPRv2 carrying either control sgRNA, sgRNA targeting *TFE3*. Subsequently, all these cells were expanded and selected with puromycin at a concentration of 2 and 5 μg mL^-1^ for 5 days. After selection, the cells were seeded in 6 cm dishes and cultured for an additional 2 days. The cells then were washed three times with fresh 75mM ammonium carbonate wash buffer (pH 7.4). Metabolites were extracted twice by adding 1.5 mL extraction solvent (40:40:20 Acetonitrile: methanol: water) at -20c for 5 minutes. The combined extracts were then centrifuged for 10min at 14000 rpm at 4c. For flow injection analysis-tandem mass spectrometry, cell extracts were diluted 1:100 in methanol. 5 μl of each sample was analyzed in two technical replicates by flow-injection coupled to an Agilent 6550 qTOF mass spectrometer (Agilent Technologies) in negative ionization mode on as previously described^109^. The mobile phase was 1 mM ammonium fluoride in isopropanol/water (60:40, v/v) and flow rate was 150 μl per min. The mobile phase was spiked with hexakis (1H,1H,3H-tetrafluoropropoxy) phosphazene and taurocholic acid ad ∼1e5 signal intensity for online mass calibration. Mass spectra were acquired in profile mode at 4 GHz (HiRes) in m/z range of 50 to 1,000. Data were collected for 0.46 minutes per sample at 1.4 spectra/s. Ions were annotated by matching their inferred mass with compounds in the HMDB database, allowing a tolerance of 1 mDa. Only deprotonated adducts were considered in the analysis.

### RNA-seq analysis

Paired-end RNA-seq reads were aligned to a Bowtie2 (v2.2.6)^110^ indexed human genome (hg38 sourced from UCSC) using STAR (2.7.1.a)^111^ with default settings. For better alignment, the first aligned splicing junctions detected by STAR (SJ.out.tab) were used for STAR alignment again with parameters “--sjdbFileChrStartEnd SJ.out.tab --readFilesIn $R1 $R2 --quantMode TranscriptoomeSAM GeneCounts --sjdbGTFfile $GTF_file --outSAMtype BAM Unsorted SortedByCoordinate”. Then the aligned transcriptome bam files (AlignedtoTranscriptome.out.bam) were used to quantify gene expression levels by rsem-calculate-expression function from RSEM(v1.3.1)^112^. Tag per million reads (TPM) from gene expression results files (.genes.results) were extracted and assembled for subsequent differential gene expression analysis by DESeq2 (1.36.0)^113^. Heatmap for deregulation of glutathione metabolism and PPP genes after ASPSCR1-TFE3 KD/KO in FU-UR-1 and sTFE were plotted in R.

### Gene Set enrichment analysis (GSEA)

GSEA was performed on expressed genes according to the software (v4.3.2) manual. GSEA was used on the Hallmark gene sets from Molecular Signatures Database (MSigDB)^114^. For the Hallmark analysis, the gene sets were ranked by the number of pairwise comparisons that had a normalized enrichment score (NES)>1 in tRCC vs other comparators. Gene sets with a nominal p value of <0.05 and an FDR of <0.25 were considered significant. FDR q value for the GSEA analysis was applied to plot bar plot of gene sets shown in the figures.

### ChIP-seq analysis

Before alignment, raw ChIP-sequencing reads were qualified by FastQC (v0.11.9). Trimmomatic (v0.39) was used to trim adaptor and low-quality reads. Trimmed reads were aligned to hg38 human genome assembly using Bowtie2 (v2.2.6)^110^, with parameters “--very-sensitive --end-to-end --mm -X 2000 --no-unal”. Proper paired and high-quality mapped reads (MAPQ >30) were extracted by samtool^115^ (v1.9) with parameter “-F 1804 -f 2 -q 30”. PCR duplicates were marked and further removed by picard tools (v2.22.3), then reads were subjected to peak calling by MACS2 (v2.1.1.20160309)^116^ with “q 0.01” for H3K27ac ChIP and “p 1e-4” for TFE3 ChIP-seq. Signal tracks for each example were generated using the MACS2 pileup function and were normalized to 1 million reads. Bigwig files were generated using the bedGraphToBigWig command for visualization. H3K27ac and TFE3 ChIP-seq signal at selected genomic loci were visualized using IGV^117^. Peak annotation was performed using annotatePeaks function from homer (v4.11.1)^118^. Enrichment analysis was performed on annotated genes proximal to TFE3 peaks using Enrichr^119^. Profile plot for TFE3 at annotated OXPHOS, glutathione metabolism and PPP genes were drawn with deeptools (v3.5.6)^120^. Bedtools (v2.29.2)^121^was used to assess overlap of TFE3 peaks among FU-UR-1, s-TFE and UOK109 cells. ROSE2^49,122^ was used to call enhancers and superenhancers based on H3K27ac signals, as well as to annotate superenhancers with nearby genes. The annotated OXPHOS genes were separated into superenhancer and typical enhancers and then their enhancer score was plotted correspondingly in R. Heatmap for enhancer score at annotated OXPHOS, TCA cycle and ETC genes were z-scored and plotted in R.

### Principal Component Analysis (PCA)

Read coverages for genomic regions of aligned H3K27ac ChIP-seq reads were computed using multiBamSummary from deeptools (v3.5.6)^120^ with default settings. PCA analysis was performed using plotPCA function from deeptools (v3.5.6)^120^ with default settings.

### GO term analysis

Gene ontology enrichment analyses were performed using EnrichR. Adjusted p values were plotted to show the significance.

### Tumor data analysis

Tumor data were obtained from TCGA cohort^3^, IMmotion151 cohort^3^ and RCC patient-derived xenograft cohorts^123^ as previously reported. For the comparison of OXPHOS, glycolysis, and *HIF1A* gene signatures between tRCC tumors and ccRCC tumors, single sample GSEA (ssGSEA) scores were computed using the GSVA package in R to infer the level of activity of OXPHOS, glycolysis, and *HIF1A* gene signatures in each cohort. In order to adjust for potential RNA-seq batch effects in visualization, signature scores were Z-scored within each dataset prior to visualization. Comparison of ssGSEA scores between tumor types was performed using Wilcoxon rank-sum tests. For the comparison of *SQSTM1* mRNA level and *HIF1A* mRNA level between tRCC tumors and ccRCC tumors, gene expression was Z-scored within each dataset independently. Comparison of gene expression scores between tumor types was performed using Wilcoxon rank-sum tests.

## Data and Materials Availability Statement

The data and unique reagents generated in this study are available upon request from the corresponding author. Analyzed data from ChIP-Seq and RNA-Seq are available in Supplementary Table S1-S3. Raw sequencing data are available in GEO under accession number GSE266517 (RNA sequencing) and GSE266530 (ChIP sequencing)

## Quantification and statistical analysis

Statistical analyses were performed by GraphPad prism 9 and Python (on Spyder v4.1.5) and R v4.3. 1. Sample sizes, statistical tests and significance are described in figure legends. Statistical comparisons were determined by Wilcoxon rank-sum test. Significance was defined as *p < 0.05, **p < 0.01, ***p < 0.001, ****p < 0.001, n.s.

## Acknowledgments

S.R.V: Doris Duke Charitable Foundation (Clinician-Scientist Development Award grant number: 2020101), Department of Defense Kidney Cancer Research Program (DoD KCRP) (W81XWH-22-1-016), Damon Runyon-Rachleff Innovation Award (Grant Number 71-22), NCI R01CA279044, V Scholar Foundation (V2022-018), Rally Foundation (23IN37). L.B-P: 1R21CA226082-01, R37CA260062. J.L.: DoD KCRP Postdoctoral and Clinical Fellowship Award (W81XWH-22-1-0399). P.K.: Kidney Cancer Association Trailblazer Award. M.G.: the Executive Committee on Research (ECOR) Fund for Medical Discovery.

## Authors’ Contributions

Conceptualization: J.L., L.B.-P., S.R.V. Performed experiments: J.L., D.S.G., M.T., P.K., B.L., C.N.W. Performed or designed analyses on genomic and/or metabolomic data: K.H, F.M., A.S., M.A., Q.X., K.H., B.A.R., M.G. Provided reagents, conceptual input, experimental design: M.G., L.B-P, S.R.V. Supervision and funding acquisition: S.R.V. Manuscript, original draft: J.L., L.B-P., S.R.V. Manuscript, editing and final draft: all authors.

## References

1. Kauffman, E. C. et al. Molecular genetics and cellular features of TFE3 and TFEB fusion kidney cancers. Nat Rev Urol 11, 465–475 (2014).

2. Moch, H., Cubilla, A. L., Humphrey, P. A., Reuter, V. E. & Ulbright, T. M. The 2016 WHO Classification of Tumours of the Urinary System and Male Genital Organs—Part A: Renal, Penile, and Testicular Tumours. European Urology 70, 93–105 (2016).

3. Bakouny, Z. et al. Integrative clinical and molecular characterization of translocation renal cell carcinoma. Cell Reports 38, 110190 (2022).

4. Argani, P. et al. Xp11 translocation renal cell carcinoma in adults: expanded clinical, pathologic, and genetic spectrum. Am J Surg Pathol 31, 1149–1160 (2007).

5. Xu, L. et al. Xp11.2 translocation renal cell carcinomas in young adults. BMC Urol 15, 57 (2015).

6. Calio, A., Segala, D., Munari, E., Brunelli, M. & Martignoni, G. MiT Family Translocation Renal Cell Carcinoma: from the Early Descriptions to the Current Knowledge. Cancers (Basel) 11, (2019).

7. Yan, X. et al. Systemic Therapy in Patients With Metastatic Xp11.2 Translocation Renal Cell Carcinoma. Clinical Genitourinary Cancer 20, 354–362 (2022).

8. Alhalabi, O. et al. Immuno-oncology (IO) combinations +/-VEGF targeted therapy (VEGF TT) in patients (pts) with advanced mit family translocation renal cell carcinomas (tRCC): Results from an international multicenter study. JCO 40, 343–343 (2022).

9. Choueiri, T. K. et al. Vascular endothelial growth factor-targeted therapy for the treatment of adult metastatic Xp11.2 translocation renal cell carcinoma. Cancer 116, 5219–5225 (2010).

10. Thouvenin, J. et al. Efficacy of Cabozantinib in Metastatic MiT Family Translocation Renal Cell Carcinomas. Oncologist oyac158 (2022) doi:10.1093/oncolo/oyac158.

11. Marcon, J. et al. Comprehensive Genomic Analysis of Translocation Renal Cell Carcinoma Reveals Copy-Number Variations as Drivers of Disease Progression. Clin. Cancer Res. (2020) doi:10.1158/1078-0432.CCR-19-3283.

12. Sun, G. et al. Integrated exome and RNA sequencing of TFE3-translocation renal cell carcinoma. Nat Commun 12, 5262 (2021).

13. Qu, Y. et al. Proteogenomic characterization of MiT family translocation renal cell carcinoma. Nat Commun 13, 7494 (2022).

14. Motzer, R. J. et al. Molecular Subsets in Renal Cancer Determine Outcome to Checkpoint and Angiogenesis Blockade. Cancer Cell 38, 803–817.e4 (2020).

15. Linehan, W. M. & Ricketts, C. J. The Cancer Genome Atlas of renal cell carcinoma: findings and clinical implications. Nat Rev Urol 16, 539–552 (2019).

16. Fukagawa, A. et al. Genomic and epigenomic integrative subtypes of renal cell carcinoma in a Japanese cohort. Nat Commun 14, 8383 (2023).

17. Riggi, N. et al. EWS-FLI1 Utilizes Divergent Chromatin Remodeling Mechanisms to Directly Activate or Repress Enhancer Elements in Ewing Sarcoma. Cancer Cell 26, 668–681 (2014).

18. Subbiah, V. & Cote, G. J. Advances in Targeting RET-Dependent Cancers. Cancer Discovery 10, 498–505 (2020).

19. Laskin, J. et al. NRG1 fusion-driven tumors: biology, detection, and the therapeutic role of afatinib and other ErbB-targeting agents. Annals of Oncology 31, 1693–1703 (2020).

20. Damayanti, N. P. et al. Therapeutic Targeting of TFE3/IRS-1/PI3K/mTOR Axis in Translocation Renal Cell Carcinoma. Clin Cancer Res 24, 5977–5989 (2018).

21. Kauffman, E. C. et al. Preclinical efficacy of dual mTORC1/2 inhibitor AZD8055 in renal cell carcinoma harboring a TFE3 gene fusion. BMC Cancer 19, 917 (2019).

22. Tsuda, M. et al. TFE3 Fusions Activate MET Signaling by Transcriptional Up-regulation, Defining Another Class of Tumors as Candidates for Therapeutic MET Inhibition. Cancer Research 67, 919–929 (2007).

23. Calcagnl, A. et al. Modelling TFE renal cell carcinoma in mice reveals a critical role of WNT signaling. Elife 5, e17047 (2016).

24. Baba, M. et al. TFE3 Xp11.2 Translocation Renal Cell Carcinoma Mouse Model Reveals Novel Therapeutic Targets and Identifies GPNMB as a Diagnostic Marker for Human Disease. Mol Cancer Res 17, 1613–1626 (2019).

25. Kobos, R. et al. Combining integrated genomics and functional genomics to dissect the biology of a cancer-associated, aberrant transcription factor, the ASPSCR1-TFE3 fusion oncoprotein. J Pathol 229, 743–754 (2013).

26. Linehan, W. M. et al. The Metabolic Basis of Kidney Cancer. Cancer Discov 9, 1006–1021 (2019).

27. Chappell, J. C., Payne, L. B. & Rathmell, W. K. Hypoxia, angiogenesis, and metabolism in the hereditary kidney cancers. Journal of Clinical Investigation 129, 442–451 (2019).

28. Kondo, K., Klco, J., Nakamura, E., Lechpammer, M. & Kaelin, W. G. Inhibition of HIF is necessary for tumor suppression by the von Hippel-Lindau protein. Cancer Cell 1, 237–246 (2002).

29. Wilde, B. R. et al. FH Variant Pathogenicity Promotes Purine Salvage Pathway Dependence in Kidney Cancer. Cancer Discovery 13, 2072–2089 (2023).

30. Gopal, R. K. et al. Early loss of mitochondrial complex I and rewiring of glutathione metabolism in renal oncocytoma. Proc. Natl. Acad. Sci. U.S.A. 115, (2018).

31. Simonnet, H. Mitochondrial complex I is deficient in renal oncocytomas. Carcinogenesis 24, 1461–1466 (2003).

32. Yong, C., Stewart, G. D. & Frezza, C. Oncometabolites in renal cancer: Warburg’s hypothesis re-examined. Nat Rev Nephrol 16, 156–172 (2020).

33. Hanahan, D. & Weinberg, R. A. The Hallmarks of Cancer. Cell 100, 57–70 (2000).

34. Ward, P. S. & Thompson, C. B. Metabolic Reprogramming: A Cancer Hallmark Even Warburg Did Not Anticipate. Cancer Cell 21, 297–308 (2012).

35. Hanahan, D. & Weinberg, R. A. Hallmarks of Cancer: The Next Generation. Cell 144, 646– 674 (2011).

36. Hanahan, D. Hallmarks of Cancer: New Dimensions. Cancer Discovery 12, 31–46 (2022).

37. Ashton, T. M., McKenna, W. G., Kunz-Schughart, L. A. & Higgins, G. S. Oxidative Phosphorylation as an Emerging Target in Cancer Therapy. Clinical Cancer Research 24, 2482–2490 (2018).

38. Yap, T. A. et al. Complex I inhibitor of oxidative phosphorylation in advanced solid tumors and acute myeloid leukemia: phase I trials. Nat Med 29, 115–126 (2023).

39. Vasan, K., Werner, M. & Chandel, N. S. Mitochondrial Metabolism as a Target for Cancer Therapy. Cell Metab 32, 341–352 (2020).

40. M, G., T, P. & L, B.-P. Reductive stress in cancer: coming out of the shadows. Trends in cancer 10, (2024).

41. Weiss-Sadan, T. et al. NRF2 activation induces NADH-reductive stress, providing a metabolic vulnerability in lung cancer. Cell Metab 35, 487–503.e7 (2023).

42. Bakouny, Z. et al. Integrative clinical and molecular characterization of translocation renal cell carcinoma. Cell Rep 38, 110190 (2022).

43. Ricketts, C. J. et al. The Cancer Genome Atlas Comprehensive Molecular Characterization of Renal Cell Carcinoma. Cell Reports 23, 313–326.e5 (2018).

44. Elias, R. et al. A renal cell carcinoma tumorgraft platform to advance precision medicine. Cell Rep 37, 110055 (2021).

45. Shuch, B., Linehan, W. M. & Srinivasan, R. Aerobic glycolysis: a novel target in kidney cancer. Expert Review of Anticancer Therapy 13, 711–719 (2013).

46. Cejas, P. et al. Subtype heterogeneity and epigenetic convergence in neuroendocrine prostate cancer. Nat Commun 12, 5775 (2021).

47. Nassar, A. H. et al. Epigenomic charting and functional annotation of risk loci in renal cell carcinoma. Nat Commun 14, 346 (2023).

48. Gopi, L. K. & Kidder, B. L. Integrative pan cancer analysis reveals epigenomic variation in cancer type and cell specific chromatin domains. Nat Commun 12, 1419 (2021).

49. Whyte, W. A. et al. Master Transcription Factors and Mediator Establish Super-Enhancers at Key Cell Identity Genes. Cell 153, 307–319 (2013).

50. Hnisz, D. et al. Super-Enhancers in the Control of Cell Identity and Disease. Cell 155, 934– 947 (2013).

51. Plitzko, B. & Loesgen, S. Measurement of Oxygen Consumption Rate (OCR) and Extracellular Acidification Rate (ECAR) in Culture Cells for Assessment of the Energy Metabolism. BIO-PROTOCOL 8, (2018).

52. Dar, S. et al. Bioenergetic Adaptations in Chemoresistant Ovarian Cancer Cells. Sci Rep 7, 8760 (2017).

53. Chen, Y. et al. Up-regulation of NMRK2 mediated by TFE3 fusions is the key for energy metabolism adaption of Xp11.2 translocation renal cell carcinoma. Cancer Lett 538, 215689 (2022).

54. Zhang, J. et al. Measuring energy metabolism in cultured cells, including human pluripotent stem cells and differentiated cells. Nat Protoc 7, 1068–1085 (2012).

55. Arroyo, J. D. et al. A Genome-wide CRISPR Death Screen Identifies Genes Essential for Oxidative Phosphorylation. Cell Metab 24, 875–885 (2016).

56. Marroquin, L. D., Hynes, J., Dykens, J. A., Jamieson, J. D. & Will, Y. Circumventing the Crabtree Effect: Replacing Media Glucose with Galactose Increases Susceptibility of HepG2 Cells to Mitochondrial Toxicants. Toxicological Sciences 97, 539–547 (2007).

57. Kierans, S. J. & Taylor, C. T. Regulation of glycolysis by the hypoxia-inducible factor (HIF): implications for cellular physiology. The Journal of Physiology 599, 23–37 (2021).

58. Tennant, D. A., Duran, R. V., Boulahbel, H. & Gottlieb, E. Metabolic transformation in cancer. Carcinogenesis 30, 1269–1280 (2009).

59. Shen, C. & Kaelin, W. G. The VHL/HIF Axis in Clear Cell Renal Carcinoma. Semin Cancer Biol 23, 18–25 (2013).

60. Brady, O. A., Martina, J. A. & Puertollano, R. Emerging roles for TFEB in the immune response and inflammation. Autophagy 14, 181–189 (2018).

61. Martina, J. A. et al. The nutrient-responsive transcription factor TFE3 promotes autophagy, lysosomal biogenesis, and clearance of cellular debris. Sci Signal 7, ra9 (2014).

62. Perera, R. M., Di Malta, C. & Ballabio, A. MiT/TFE Family of Transcription Factors, Lysosomes, and Cancer. Annu Rev Cancer Biol 3, 203–222 (2019).

63. Keshet, R., Szlosarek, P., Carracedo, A. & Erez, A. Rewiring urea cycle metabolism in cancer to support anabolism. Nat Rev Cancer 18, 634–645 (2018).

64. Chakraborty, S., Balan, M., Sabarwal, A., Choueiri, T. K. & Pal, S. Metabolic reprogramming in renal cancer: Events of a metabolic disease. Biochimica et Biophysica Acta (BBA) - Reviews on Cancer 1876, 188559 (2021).

65. Hajaj, E., Sciacovelli, M., Frezza, C. & Erez, A. The context-specific roles of urea cycle enzymes in tumorigenesis. Molecular Cell 81, 3749–3759 (2021).

66. Khare, S. et al. ASS1 and ASL suppress growth in clear cell renal cell carcinoma via altered nitrogen metabolism. Cancer & Metabolism 9, 40 (2021).

67. Lash, L. H. Role of glutathione transport processes in kidney function. Toxicology and Applied Pharmacology 204, 329–342 (2005).

68. Priolo, C. et al. Impairment of gamma-glutamyl transferase 1 activity in the metabolic pathogenesis of chromophobe renal cell carcinoma. Proc. Natl. Acad. Sci. U.S.A. 115, (2018).

69. Xiao & Meierhofer. Glutathione Metabolism in Renal Cell Carcinoma Progression and Implications for Therapies. IJMS 20, 3672 (2019).

70. Pillai, R., Hayashi, M., Zavitsanou, A.-M. & Papagiannakopoulos, T. NRF2: KEAPing Tumors Protected. Cancer Discovery 12, 625–643 (2022).

71. Romero, R. et al. Keap1 mutation renders lung adenocarcinomas dependent on Slc33a1. Nat Cancer 1, 589–602 (2020).

72. Xiao, W. & Loscalzo, J. Metabolic Responses to Reductive Stress. Antioxidants & Redox Signaling 32, 1330–1347 (2020).

73. Zhao, Y. et al. SoNar, a Highly Responsive NAD+/NADH Sensor, Allows High-Throughput Metabolic Screening of Anti-tumor Agents. Cell Metabolism 21, 777–789 (2015).

74. Titov, D. V. et al. Complementation of mitochondrial electron transport chain by manipulation of the NAD ^+^ /NADH ratio. Science 352, 231–235 (2016).

75. Dempster, J. M. et al. Chronos: a cell population dynamics model of CRISPR experiments that improves inference of gene fitness effects. Genome Biol 22, 343 (2021).

76. Dempster, J. M. et al. Agreement between two large pan-cancer CRISPR-Cas9 gene dependency data sets. Nat Commun 10, 5817 (2019).

77. Wolf, M. M., Rathmell, W. K. & Beckermann, K. E. Modeling Clear Cell Renal Cell Carcinoma and Therapeutic Implications. Oncogene 39, 3413–3426 (2020).

78. Molina, J. R. et al. An inhibitor of oxidative phosphorylation exploits cancer vulnerability. Nat Med 24, 1036–1046 (2018).

79. Birsoy, K. et al. An Essential Role of the Mitochondrial Electron Transport Chain in Cell Proliferation Is to Enable Aspartate Synthesis. Cell 162, 540–551 (2015).

80. Ivan, M. & Kaelin, W. G. The EGLN-HIF O 2 -Sensing System: Multiple Inputs and Feedbacks. Molecular Cell 66, 772–779 (2017).

81. Ivan, M. et al. HIFalpha targeted for VHL-mediated destruction by proline hydroxylation: implications for O2 sensing. Science 292, 464–468 (2001).

82. Shen, C. et al. Genetic and functional studies implicate HIF1a as a 14q kidney cancer suppressor gene. Cancer Discov 1, 222–235 (2011).

83. Kim, J., Tchernyshyov, I., Semenza, G. L. & Dang, C. V. HIF-1-mediated expression of pyruvate dehydrogenase kinase: A metabolic switch required for cellular adaptation to hypoxia. Cell Metabolism 3, 177–185 (2006).

84. Miska, J. et al. HIF-1a Is a Metabolic Switch between Glycolytic-Driven Migration and Oxidative Phosphorylation-Driven Immunosuppression of Tregs in Glioblastoma. Cell Reports 27, 226–237.e4 (2019).

85. Chen, N. et al. Roxadustat for Anemia in Patients with Kidney Disease Not Receiving Dialysis. N Engl J Med 381, 1001–1010 (2019).

86. Mittelman, M. et al. Efficacy and Safety of Roxadustat for Treatment of Anemia in Patients with Lower-Risk Myelodysplastic Syndrome (LR-MDS) with Low Red Blood Cell (RBC) Transfusion Burden: Results of Phase III Matterhorn Study. Blood 142, 195–195 (2023).

87. Malouf, G. G. et al. Next-Generation Sequencing of Translocation Renal Cell Carcinoma Reveals Novel RNA Splicing Partners and Frequent Mutations of Chromatin-Remodeling Genes. Clin Cancer Res 20, 4129–4140 (2014).

88. Bezwada, D. et al. Mitochondrial metabolism in primary and metastatic human kidney cancers. *bioRxiv* 2023.02.06.527285 (2023) doi:10.1101/2023.02.06.527285.

89. Monda, S. M. et al. The Metastatic Risk of Renal Cell Carcinoma by Primary Tumor Size and Subtype. European Urology Open Science 52, 137–144 (2023).

90. Mir, M. C. et al. Altered transcription factor E3 expression in unclassified adult renal cell carcinoma indicates adverse pathological features and poor outcome. BJU International 108, (2011).

91. Sicinska, E. et al. ASPSCR1::TFE3 Drives Alveolar Soft Part Sarcoma by Inducing Targetable Transcriptional Programs. Cancer Research (2024) doi:10.1158/0008-5472.CAN-23-2115.

92. Pastore, N. et al. TFE3 regulates whole-body energy metabolism in cooperation with TFEB. EMBO Mol Med 9, 605–621 (2017).

93. Zhou, Y. et al. Recent advances of mitochondrial complex I inhibitors for cancer therapy: Current status and future perspectives. European Journal of Medicinal Chemistry 251, 115219 (2023).

94. Chandel, N. Metformin Inhibits Mitochondrial Complex I To Promote Health. Innovation in Aging 5, 454 (2021).

95. Wheaton, W. W. et al. Metformin inhibits mitochondrial complex I of cancer cells to reduce tumorigenesis. eLife 3, e02242 (2014).

96. Price, C. et al. Genome-Wide Interrogation of Human Cancers Identifies EGLN1 Dependency in Clear Cell Ovarian Cancers. Cancer Res 79, 2564–2579 (2019).

97. Singh, A. K. et al. Daprodustat for the Treatment of Anemia in Patients Not Undergoing Dialysis. N Engl J Med 385, 2313–2324 (2021).

98. Chertow, G. M. et al. Vadadustat in Patients with Anemia and Non–Dialysis-Dependent CKD. N Engl J Med 384, 1589–1600 (2021).

99. Jang, S. et al. Elucidating the contribution of mitochondrial glutathione to ferroptosis in cardiomyocytes. Redox Biology 45, 102021 (2021).

100. Takahashi, N. et al. 3D Culture Models with CRISPR Screens Reveal Hyperactive NRF2 as a Prerequisite for Spheroid Formation via Regulation of Proliferation and Ferroptosis. Mol Cell 80, 828–844.e6 (2020).

101. Clerici, S. & Boletta, A. Role of the KEAP1-NRF2 Axis in Renal Cell Carcinoma. Cancers 12, 3458 (2020).

102. Cancer Genome Atlas Research Network et al. Comprehensive Molecular Characterization of Papillary Renal-Cell Carcinoma. N Engl J Med 374, 135–145 (2016).

103. Li, L. et al. SQSTM1 Is a Pathogenic Target of 5q Copy Number Gains in Kidney Cancer. Cancer Cell 24, 738–750 (2013).

104. Sayin, V. I. et al. Activation of the NRF2 antioxidant program generates an imbalance in central carbon metabolism in cancer. eLife 6, e28083 (2017).

105. Viswanathan, S. R. et al. Genome-scale analysis identifies paralog lethality as a vulnerability of chromosome 1p loss in cancer. Nat Genet 50, 937–943 (2018).

106. Li, J. et al. An alternative CTCF isoform antagonizes canonical CTCF occupancy and changes chromatin architecture to promote apoptosis. Nat Commun 10, 1535 (2019).

107. Stringer, C., Wang, T., Michaelos, M. & Pachitariu, M. Cellpose: a generalist algorithm for cellular segmentation. Nat Methods 18, 100–106 (2021).

108. Stirling, D. R. et al. CellProfiler 4: improvements in speed, utility and usability. BMC Bioinformatics 22, 433 (2021).

109. Fuhrer, T., Heer, D., Begemann, B. & Zamboni, N. High-throughput, accurate mass metabolome profiling of cellular extracts by flow injection-time-of-flight mass spectrometry. Anal Chem 83, 7074–7080 (2011).

110. Langmead, B. & Salzberg, S. L. Fast gapped-read alignment with Bowtie 2. Nat Methods 9, 357–359 (2012).

111. Dobin, A. et al. STAR: ultrafast universal RNA-seq aligner. Bioinformatics 29, 15–21 (2013).

112. Li, B. & Dewey, C. N. RSEM: accurate transcript quantification from RNA-Seq data with or without a reference genome. BMC Bioinformatics 12, 323 (2011).

113. Love, M. I., Huber, W. & Anders, S. Moderated estimation of fold change and dispersion for RNA-seq data with DESeq2. Genome Biol 15, 550 (2014).

114. Liberzon, A. et al. The Molecular Signatures Database (MSigDB) hallmark gene set collection. Cell Syst 1, 417–425 (2015).

115. Li, H. et al. The Sequence Alignment/Map format and SAMtools. Bioinformatics 25, 2078– 2079 (2009).

116. Zhang, Y. et al. Model-based analysis of ChIP-Seq (MACS). Genome Biol 9, R137 (2008).

117. Robinson, J. T. et al. Integrative genomics viewer. Nat Biotechnol 29, 24–26 (2011).

118. Heinz, S. et al. Simple combinations of lineage-determining transcription factors prime cis-regulatory elements required for macrophage and B cell identities. Mol Cell 38, 576–589 (2010).

119. Chen, E. Y. et al. Enrichr: interactive and collaborative HTML5 gene list enrichment analysis tool. BMC Bioinformatics 14, 128 (2013).

120. Ramfrez, F. et al. deepTools2: a next generation web server for deep-sequencing data analysis. Nucleic Acids Res 44, W160–165 (2016).

121. Quinlan, A. R. & Hall, I. M. BEDTools: a flexible suite of utilities for comparing genomic features. Bioinformatics 26, 841–842 (2010).

122. Loven, J. et al. Selective inhibition of tumor oncogenes by disruption of super-enhancers. Cell 153, 320–334 (2013).

123. Elias, R. et al. A renal cell carcinoma tumorgraft platform to advance precision medicine. Cell Rep 37, 110055 (2021).

124. Gopi, L. K. & Kidder, B. L. Integrative pan cancer analysis reveals epigenomic variation in cancer type and cell specific chromatin domains. Nat Commun 12, 1419 (2021).

125. Li, L. et al. SQSTM1 Is a Pathogenic Target of 5q Copy Number Gains in Kidney Cancer. Cancer Cell 24, 738–750 (2013).

