## Supplementary Figures and Captions for "TFE3 fusions direct an oncogenic transcriptional program that drives OXPHOS and unveils vulnerabilities in translocation renal cell carcinoma"

### Extended Data Figures

### Extended Data Fig. S1: tRCCs display activation of OXPHOS programs.

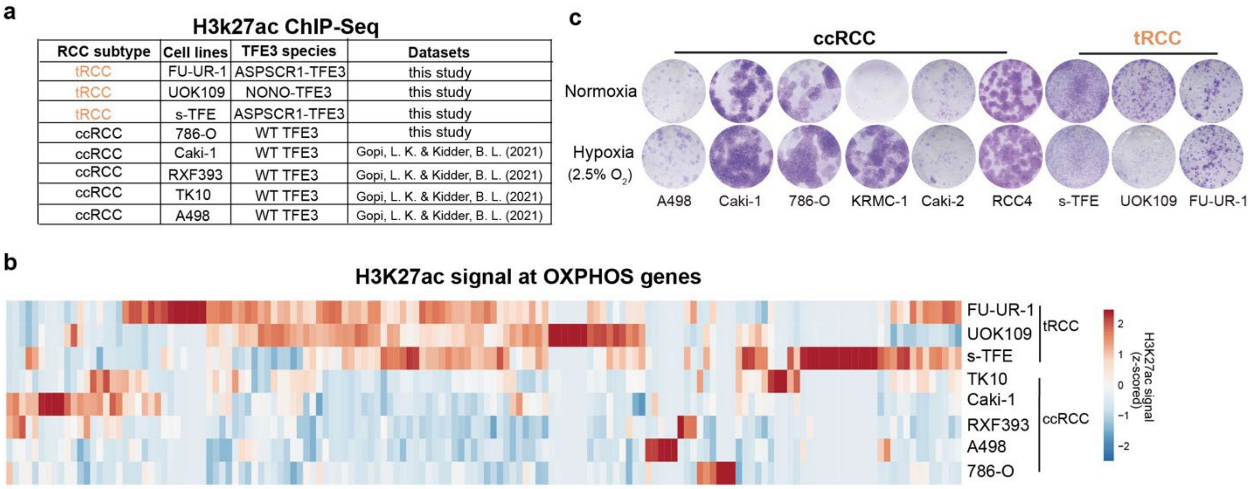


**Extended Data Fig. S1**

(a) Source of H3K27ac ChIP-Seq data from ccRCC or tRCC cell lines analyzed in this study.

(b) H3K27ac signal (quantified by ROSE2, Methods) at enhancers in proximity to OXPHOS genes in ccRCC and tRCC cell lines.

(c) Crystal violet assay of ccRCC and tRCC cultured under normoxic or hypoxic conditions.

### Extended Data Fig. S2: Transcriptional activation of OXPHOS genes by TFE3 fusions.

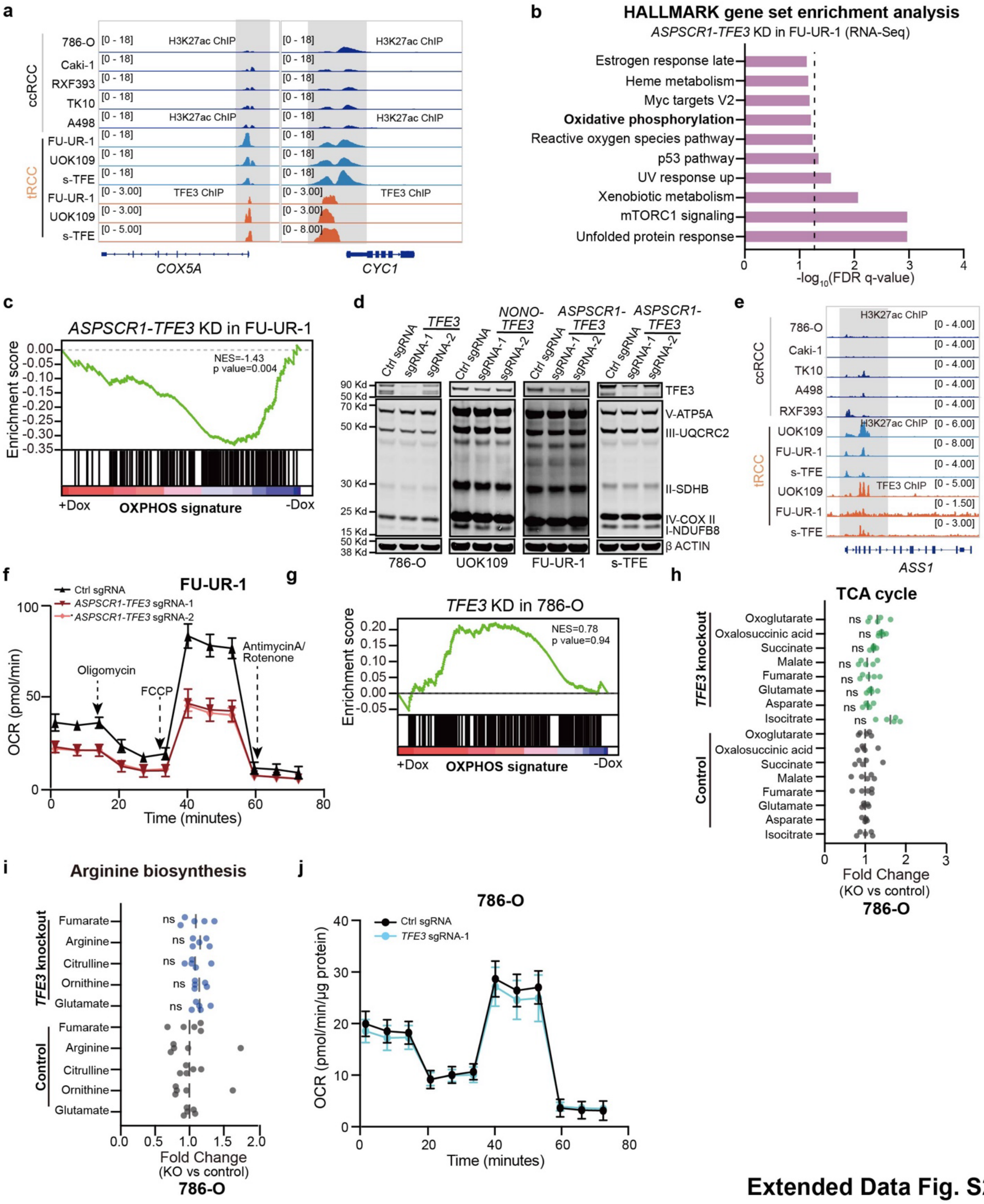


**Extended Data Fig. S2**

(a) IGV snapshot showing TFE3 fusion (orange track) and H3K27ac (light or dark blue tracks) signal at representative OXPHOS related loci in ccRCC and tRCC cell lines.

(b) Bar plot showing the top gene sets depleted upon *ASPSCR1-TFE3* knockout in FU-UR-1 cells.

(c) GSEA plot showing depletion of OXPHOS gene signature in FU-UR-1 cells upon *ASPSCR1-TFE3* knockout.

(d) Western blot for TFE3, OXPHOS after knockout of *TFE3* or *TFE3* fusion in ccRCC cell line (786-O) or tRCC cell lines (UOK109, FU-UR-1, s-TFE).

(e) IGV snapshot showing TFE3 fusion (orange track) and H3K27ac (light or dark blue tracks) signal at *ASS1* locus in ccRCC and tRCC cell lines.

(f) OCR after knockdown of *ASPSCR1-TFE3* in FU-UR-1cell line. Data shown as mean ± s.d, n=5-6 biological replicates.

(g) GSEA plot showing no significant change of OXPHOS gene signature in 786-O ccRCC cells upon *TFE3* knockdown.

(h) Change in levels of TCA cycle-related metabolites following *TFE3* knockout in 786-O cells. For each metabolite, fold change was normalized to control sgRNA condition. Data are shown as mean ± s.d, n=5 biological replicates per cell line.

(i) Change in levels of arginine biosynthesis-related metabolites following *TFE3* knockout in 786-O cells. For each metabolite, fold change was normalized to control sgRNA condition. Data are shown as mean ± s.d, n=5 biological replicates per cell line.

(j) OCR after knockout of *TFE3* in 786-O ccRCC cell line. Data are shown as mean ± s.d, n=5-6 biological replicates.

For panels (h-i), statistical significance was determined by Mann-Whitney U test. *$p < 0.05$, **$p < 0.01$, ***$p < 0.001$, **** $p < 0.0001$, n.s. not significant.

### Extended Data Fig. S3: Regulation of redox balance by TFE3 fusions in tRCC

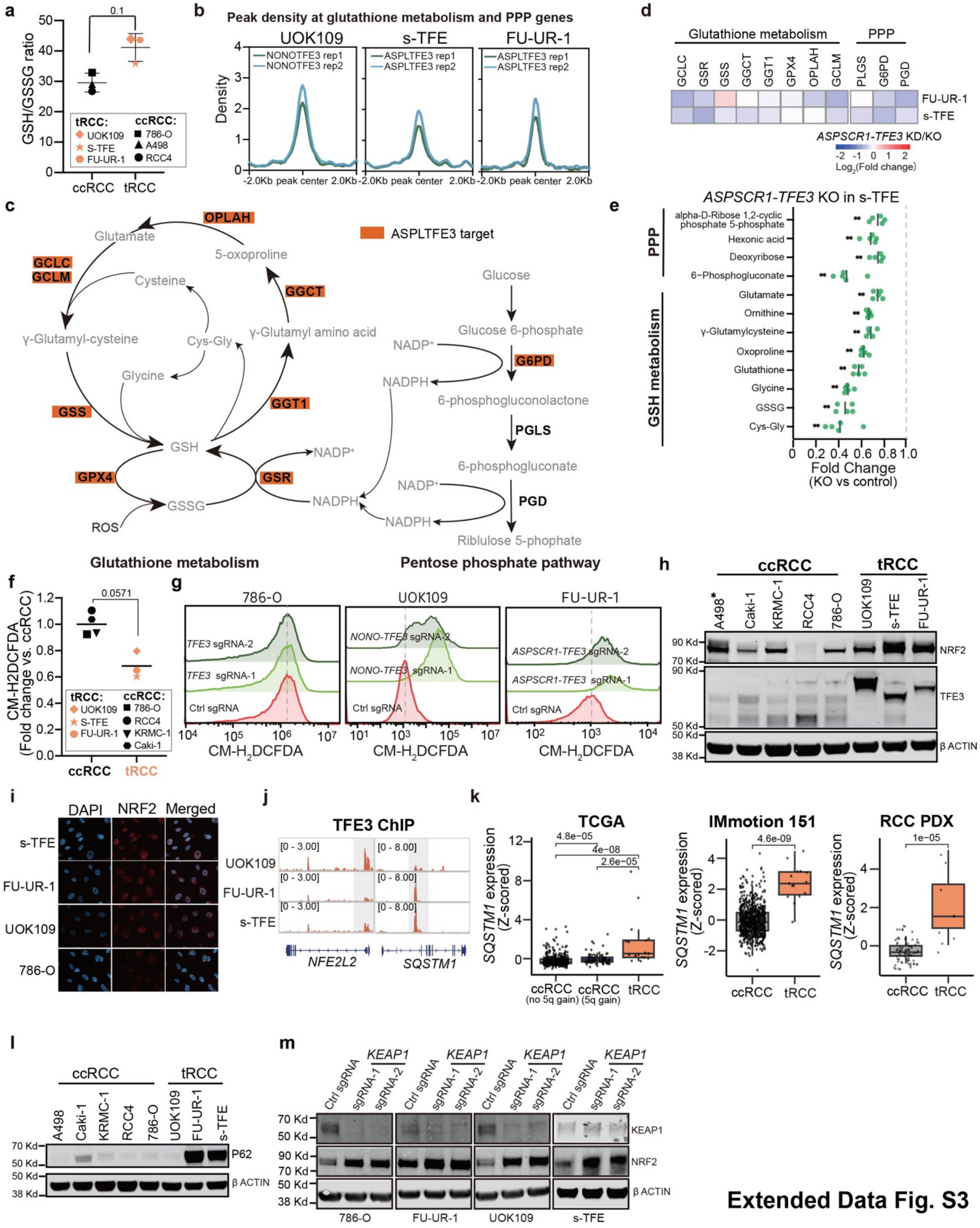


**Extended Data Fig. S3**

(a) Ratio of GSH to GSSG levels as measured by untargeted metabolomics in three ccRCC cell lines (786-O, A498, RCC4) vs. three tRCC cell lines (UOK109, FU-UR-1, s-TFE). Data are shown as mean ± s.d, n=5 biological replicates per cell line.

(b) Profile plot showing TFE3 fusion ChIP-Seq signal at glutathione metabolism and pentose phosphate pathway (PPP) genes.

(c) Schematic of glutathione metabolism and PPP, annotated with genes that are ASPL-TFE3 targets as determined by ChIP-Seq in s-TFE cells (orange boxes). In the schematic, enzymes are in black text, metabolites are in gray text.

(d) Heatmap showing the change in expression of glutathione metabolism and PPP-related genes following *ASPSCR1-TFE3* knockout in s-TFE cells or knockdown in FU-UR-1 cells.

(e) Change in levels of glutathione metabolism and PPP-related metabolites following *ASPSCR1-TFE3* knockout in s-TFE cells. For each metabolite, fold change was normalized to control sgRNA condition. Data are shown as mean ± s.d, n=5 biological replicates per cell line.

(f) Intracellular ROS level in ccRCC and tRCC cell lines. Quantification of intracellular ROS levels via CM-$H_2$DCFDA indicator in tRCC (n=3, UOK109, FU-UR-1, s-TFE) and ccRCC cell lines (n=5, 786-O, Caki-1, KRMC-1, RCC4, A498).

(g) Intracellular ROS levels (measured by CM-$H_2$DCFDA indicator) after *TFE3* knockout in ccRCC cell line (786-O) and tRCC cell lines (UOK109, FU-UR-1).

(h) Western blot showing TFE3 and NRF2 protein levels across a panel of ccRCC (n=5) and tRCC (n=3) cell lines. Note, A498 (ccRCC) cells have been previously reported to have high NRF2 activation, possibly due to *SQSTM1* overexpression and/or *CUL3* mutation[125].

(i) Immunofluorescence showing subcellular localization of NRF2 in tRCC cell lines vs. a ccRCC cell line (786-O).

(j) IGV snapshot showing TFE3 fusion signal at *NFE2L2* and *SQSTM1* loci in tRCC cell lines (UOK109, FU-UR-1, s-TFE).

(k) *SQSTM1* mRNA level in ccRCC or tRCC tumors from three independent studies (TCGA, Motzer et al., Elias et al. (PDX)).

(l) Western blot of p62 expression in ccRCC (n=5, A498, Caki-1, KRMC-1, RCC4, 786-O) and tRCC (n=3, UOK109, FU-UR-1, s-TFE) cell lines.

(m) Western blot showing NRF2 and KEAP1 levels following *KEAP1* knockout in ccRCC cell line (786-O) and tRCC cell lines (UOK109, FU-UR-1, s-TFE).

For panel (a), (e-f) and (k), statistical significance was determined by Mann-Whitney U test. *$p < 0.05$, **$p < 0.01$, ***$p < 0.001$, **** $p < 0.0001$, n.s. not significant.

**Extended Data Fig. S4: Induction of reductive stress by NRF2 activation in tRCC cells.**

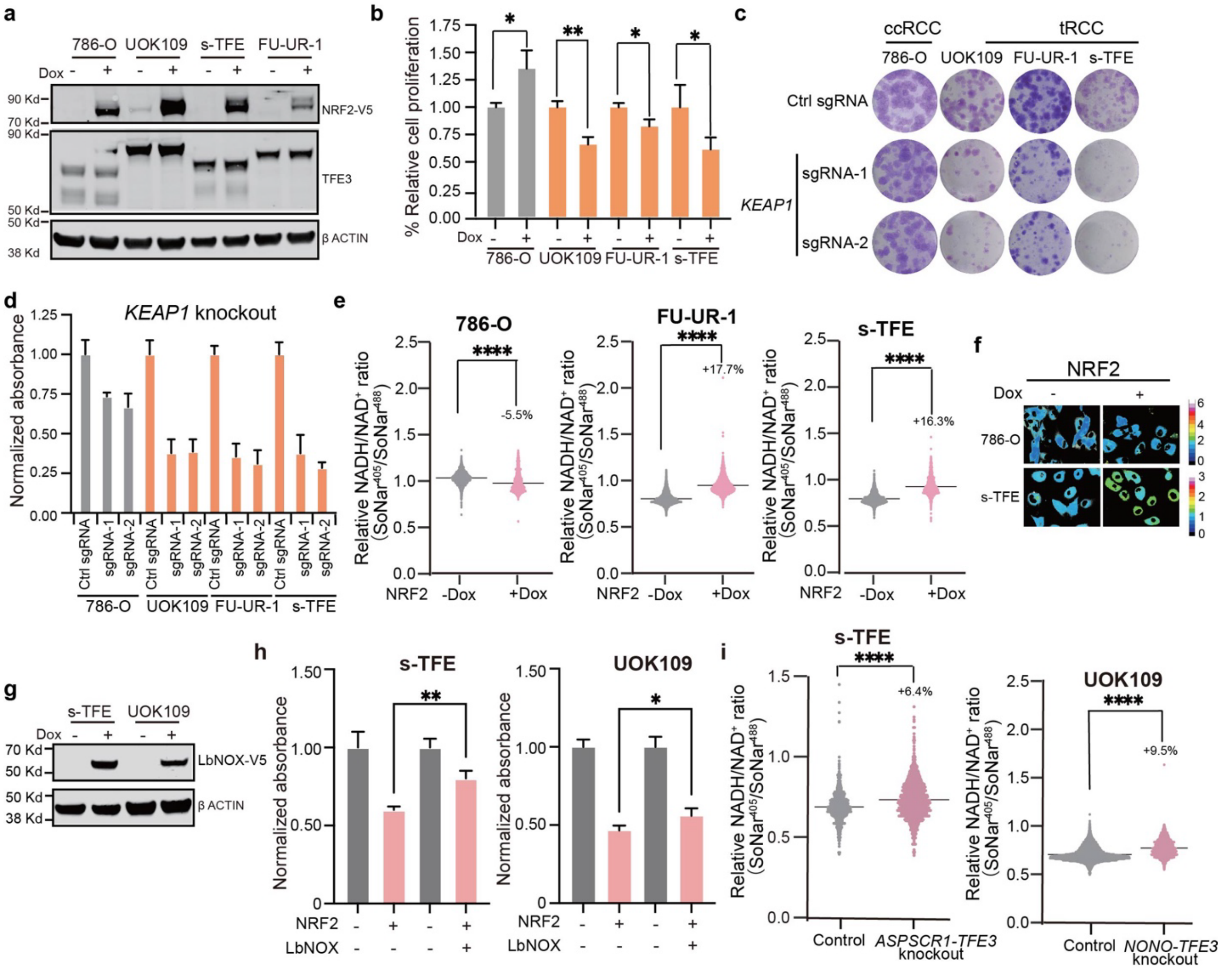


**Extended Data Fig. S4**

(a) Western blot showing doxycycline-inducible V5-tagged NRF2 overexpression in ccRCC (786-O) and tRCC cells (UOK109, FU-UR-1, s-TFE), detected by V5 antibody.

(b) Quantification of clonogenic capacity (Crystal Violet assay) after knockout of *KEAP1* in ccRCC (786-O) or tRCC (UOK109, FU-UR-1, s-TFE) cell lines. Data are shown as mean ± s.d, n=3 biological replicates.

(c) Representative well for colony formation in ccRCC and tRCC cells upon *KEAP1* knockout.

(d) Quantification of clonogenic capacity (Crystal Violet assay) after knockout of *KEAP1* in ccRCC (786-O) or tRCC (UOK109, FU-UR-1, s-TFE) cell lines. Data are shown as mean ± s.d, n=3 biological replicates.

(e) Quantification of NADH to NAD+ ratio via SoNar assay following NRF2 overexpression in ccRCC cell line(786-O) and tRCC cell lines (FU-UR-1 and s-TFE).

(f) Representative images displaying NADH/NAD+ ratio as assessed by SoNar assay in tRCC vs. ccRCC cells (images pseudocolored by NADH/$NAD^+$ ratio).

(g) Western blot confirming LbNOX overexpression in UOK109 and s-TFE cells transduced with lentiviral vector encoding doxycycline-inducible *LbNOX-V5*.

(h) Quantification of clonogenic capacity (Crystal Violet assay) after doxycycline-inducible expression of NRF2, with or without co-expression of the NADH oxidizing enzyme LbNOX in UOK109 and s-TFE cells. Data are shown as mean ± s.d, n=3 biological replicates.

(i) Quantification of NADH to NAD+ ratio via SoNar assay after *TFE3* fusion knockout in tRCC cell lines (UOK109 and s-TFE).

For panels (b), (e), (h) and (i), statistical significance was determined by Mann-Whitney U test. *p < 0.05, **p < 0.01, ***p < 0.001, **** p < 0.0001, n.s. not significant.

### Extended Data Fig. S5: *EGLN1* and *VHL* are selectively essential in tRCC cells

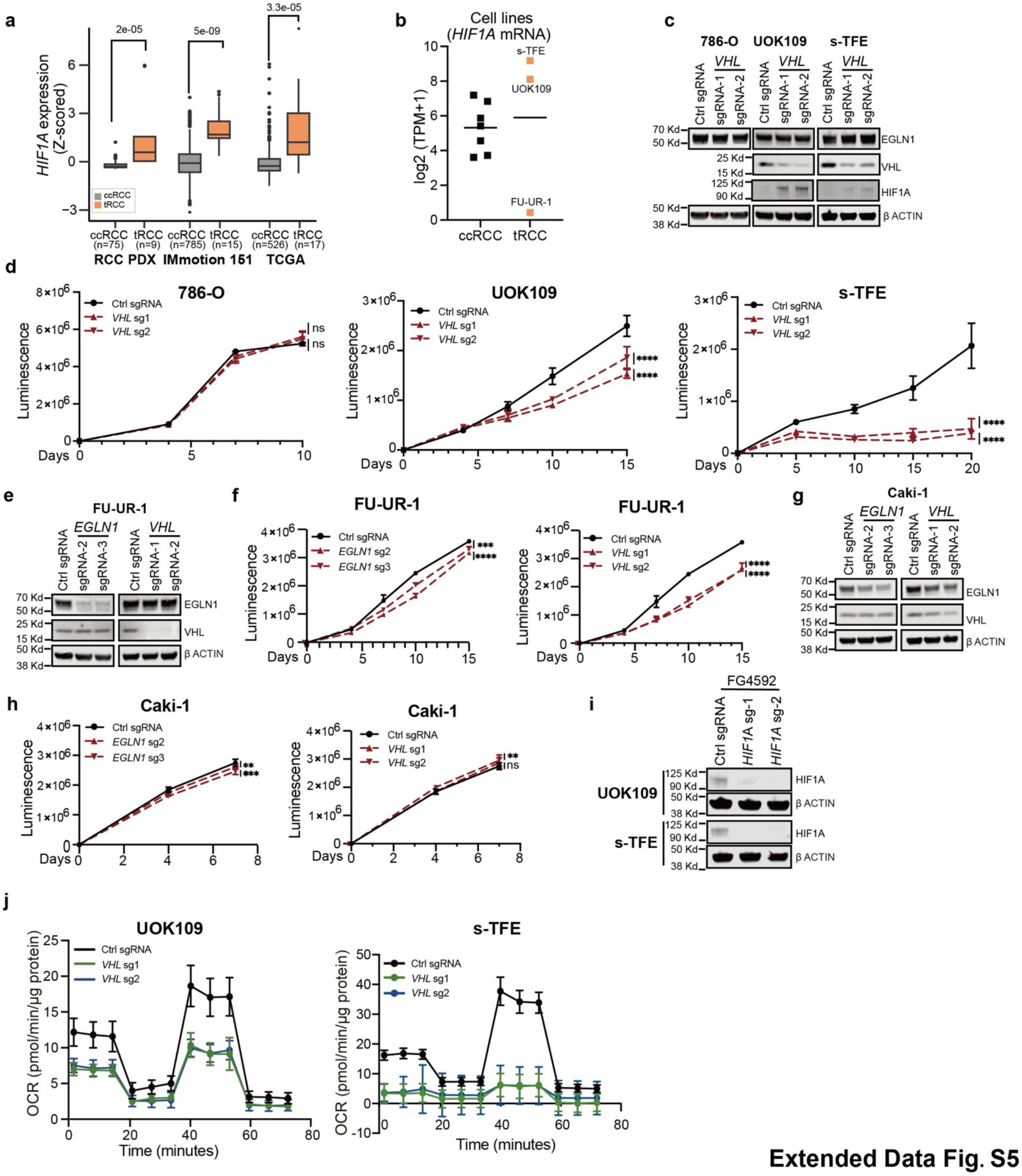


**Extended Data Fig. S5**

(a) *HIF1A* mRNA level in ccRCC or tRCC tumors from three independent studies (TCGA, Motzer et al., Elias et al. (PDX)).

(b) *HIF1A* mRNA level in tRCC cell lines (n=3, UOK109, FU-UR-1, s-TFE) versus ccRCC cell lines (n=7, A498, A704, 786-O, 769-P, Caki-1, Caki-2, OS-RC-2).

(c) Western blot showing the expression of EGLN1, VHL, HIF1A after knockout of *VHL* in ccRCC (786-O) and tRCC (UOK109, sTFE and FU-UR-1) cell lines.

(d) Cell proliferation of tRCC cell lines (UOK109 and s-TFE) and ccRCC (786-O) cell lines after knockout of *VHL*. Data are shown as mean ± s.d. for n= 3 biological replicates.

(e) Western blot showing the expression of EGLN1, VHL after knockout of *EGLN1* or *VHL* in ccRCC (FU-UR-1) cell line.

(f) Cell proliferation of FU-UR-1 cells after knockout of *EGLN1* or *VHL*. Data are shown as mean ± s.d. for n= 3 biological replicates.

(g) Western blot showing the expression of EGLN1, VHL after knockout of *EGLN1* or *VHL* in tRCC (Caki-1) cell line.

(h) Cell proliferation of Caki-1 cells (*VHL* preserved) after knockout of *EGLN1* or *VHL*. Data are shown as mean ± s.d. for n= 3 biological replicates.

(i) Western blot showing the HIF1A expression in UOK109 and s-TFE expressing sgRNA target *HIF1A* after treatment with EGLN1 inhibitor (FG4592).

(j) OCR after knockout of *VHL* in UOK109 and s-TFE cell lines. Data are shown as mean ± s.d, n=8-14 biological replicates.

For panels (a), (d), (f) and (h), statistical significance was determined by Mann-Whitney U test. *$p < 0.05$, **$p < 0.01$, ***$p < 0.001$, **** $p < 0.0001$, n.s. not significant.
